# The BLM DHBN domain and structure-selective nucleases drive mitotic arrest-dependent telomere deprotection under WRN and TRF2 control

**DOI:** 10.64898/2026.08.19.745663

**Authors:** Placide Niyonshuti, Makoto T. Hayashi

## Abstract

Telomeres shield chromosome ends from DNA damage response through T-loops, lariat DNA structures formed and stabilized by the shelterin protein TRF2. During prolonged mitotic arrest, telomeres lose this protection through a process termed mitotic arrest-dependent (MAD) telomere deprotection, which elicits telomere-specific DNA damage signaling in the absence of telomere shortening or chromosome end-to-end fusions. We previously demonstrated that the RecQ helicase BLM promotes MAD telomere deprotection, whereas the related helicase WRN suppresses it independently of its catalytic activities. However, the molecular interplay between BLM and WRN at mitotic telomeres and whether additional recombination-associated enzymes contribute to MAD telomere deprotection have remained unresolved. Here, we identify the Dimerization Helical Bundle in the N-terminal (DHBN) domain of BLM helicase as the critical determinant of MAD telomere deprotection and show that WRN selectively restrains this activity without interfering with BLM’s canonical genome-protective functions. We further show that both MUS81 and GEN1 contribute to MAD telomere deprotection. Moreover, the exacerbation of MAD telomere deprotection observed upon TRF2 depletion is strongly attenuated by additional depletion of these enzymes, demonstrating that TRF2 normally protects T-loop junctions from their enzymatic activities. Collectively, our findings reveal how telomeres become selectively vulnerable during prolonged mitotic arrest and uncover a regulated enzymatic mechanism that repurposes recombination machinery at chromosome ends.

## Introduction

Telomeres are specialized nucleoprotein structures that protect eukaryotic chromosome ends from being recognized as DNA double-strand breaks. In mammalian somatic cells, this protection is achieved through the coordinated action of the shelterin complex members, particularly TRF2, which both suppresses DNA damage signaling [1, 2] and promotes the formation and maintenance of a lariat-like DNA architecture known as the telomere loop (T-loop) [3, 4]. The T-loop is proposed to form when the 3′ single-stranded telomeric overhang invades duplex telomeric DNA, creating a displacement loop (D-loop) [5, 6]. Telomeres harboring intact T-loops are protected from nucleolytic attack and helicase activity that would otherwise expose chromosome termini, thereby preventing DDR activation and chromosome end-to-end fusions [3, 4, 7]. The loss of TRF2 and other shelterin complex members, together with progressive shortening of telomeric DNA due to the end-replication problem, is widely accepted as the principal trigger of telomere DDR [8, 9]. In addition, accumulating evidence indicates that T-loop functions in somatic cells can be compromised under conditions of prolonged mitotic arrest [7, 10–17].

Mitotic chromosomes are highly condensed, DNA repair pathways are globally attenuated, and telomeres remain protected despite extensive structural reorganization during normal mitosis [18]. Previous work has established that extended mitotic arrest leads to a phenomenon termed mitotic arrest-dependent (MAD) telomere deprotection (also referred to as mitotic telomere deprotection), characterized by the appearance of telomere deprotection-induced foci (TIF) in the absence of telomere shortening or overt chromosome end-to-end fusions [10]. Consistent with active T-loop resolution during mitotic arrest, linearization of purified mitotic telomere molecules has been reported [7]. Our conceptual framework for understanding MAD telomere deprotection is grounded in the structural similarities between telomeric loops and homologous recombination (HR) intermediates [17]. T-loops share key features with D-loops and potentially with single-and double-Holliday junctions (dHJs), including strand invasion, heteroduplex DNA formation, and topological constraints amenable to active resolution [6, 17]. In the context of HR, these structures are processed by multiple enzymes, including the Bloom syndrome helicase (BLM), the crossover junction endonuclease MUS81, and the structure-selective endonuclease GEN1 [19]. BLM belongs to the RecQ helicase family and serves as a central regulator of genome stability, functioning primarily within the BTRR (BLM-TOP3A-RMI1-RMI2) complex to suppress sister chromatid exchange (SCE) by dissolving dHJs [20]. Operating during S and G2 phases, BLM promotes high-fidelity repair by facilitating DNA end resection while concurrently disrupting aberrant RAD51 filaments to prevent illegitimate recombination [21, 22]. A defining structural feature of BLM is the Dimerization Helical Bundle in the N-terminal (DHBN) domain, which mediates protein oligomerization and efficient dissolution of dHJs [20].

We have recently reported that MAD telomere deprotection requires BLM’s helicase activity, its interaction with other BTRR components, TOP3A enzymatic activity, and all remaining members of the BTRR complex, collectively suggesting that the BTRR-mediated enzymatic activity drives dissolution of the mitotic T-loop junction [16]. We have also demonstrated that the RecQ helicase WRN antagonizes MAD telomere deprotection through a non-enzymatic mechanism and have identified a discrete WRN domain, comprising coiled-coil-forming residues 168-333, that is sufficient for this mechanism [15]. Nevertheless, the underlying molecular mechanism of WRN-dependent suppression remains incompletely understood. Furthermore, the exposure of the T-loop junction to the BTRR complex during mitotic arrest raises the possibility that the Holliday junction resolvases MUS81 and GEN1 also contribute to MAD telomere deprotection.

In this study, we combined genetic, biochemical, and live-cell imaging approaches to elucidate the molecular mechanisms underlying MAD telomere deprotection. We systematically interrogated the functional interplay between BLM and WRN and investigated the contributions of MUS81 and GEN1 to MAD telomere deprotection. Furthermore, we examined how these enzymatic activities are regulated by TRF2-mediated telomere chromatin organization. Our findings support a model in which MAD telomere deprotection results from the active processing of T-loop structures by recombination-associated enzymes, a process mechanistically distinct from canonical sister chromatid-based recombination. Specifically, we demonstrate that the BLM DHBN domain is essential for promoting MAD telomere deprotection and is targeted by the suppressive WRN coiled-coil domain, establishing it as a regulatory hub for this process. We further show that the catalytic activities of GEN1 and MUS81 contribute to MAD telomere deprotection. Moreover, depletion of these enzymes markedly attenuates the elevated MAD telomere deprotection observed upon TRF2 loss, demonstrating that TRF2 normally suppresses these enzymatic activities at mitotic telomeres. Finally, we demonstrate that the interaction between the TRF2 basic domain and core histones is destabilized during mitotic arrest, suggesting that modulation of the TRF2 basic domain–histone interaction during mitotic arrest may contribute to the licensing of helicase and resolvase activities at the T-loop.

## Result

### The BLM DHBN domain is required for MAD telomere deprotection

BLM promotes MAD telomere deprotection [16], whereas WRN overexpression suppresses it via a non-enzymatic mechanism dependent on the coiled-coil region (CCR) [15]. We hypothesized that WRN exerts its suppressive effect by physically interacting with factors involved in MAD telomere deprotection. In silico screening using Predictome [23], a structure-based protein–protein interaction prediction tool, suggested that the CCR of WRN potentially interacts with the BLM DHBN domain, which is required for BLM homodimerization [20, 24]. Notably, the presence of WRN was predicted to reduce the predicted local distance difference test (pLDDT) scores of the BLM DHBN domain, suggesting that the DHBN adopts a locally disordered conformation within the WRN–BLM heterodimer (Supplementary Figure 1a).

To validate this predicted protein–protein interaction, we performed co-immunoprecipitation (Co-IP) assays in IMR-90 hTERT E6E7 cells (immortalized by hTERT and transformed by the HPV16-derived oncoproteins E6 and E7) co-expressing Flag-tagged wild-type WRN together with one of three BLM variants: wild-type BLM (BLM^WT^), a DHBN domain deletion mutant (BLM^ΔDHBN^), or a BLM variant harboring point mutations in a cluster of hydrophobic residues within the DHBN (BLM^Patch-8A^) [20] (Figure 1a). BLM^Patch-8A^ has previously been shown to lack dimerization capacity and full anti-SCE activity [20]. Co-IP analysis revealed that wild-type WRN and BLM interact under basal conditions, as evidenced by robust co-precipitation of BLM from Flag-WRN^WT^ immunoprecipitates (Figure 1b and Supplementary Figure 1b). This interaction was markedly diminished in cells expressing BLM^ΔDHBN^ but remained largely intact in cells expressing BLM^Patch-8A^ (Figure 1b and Supplementary Figure 1b). Furthermore, treatment with colcemid for 24 hours to induce a prolonged mitotic arrest attenuated this interaction (Figure 1b and Supplementary Figure 1b). Collectively, these results demonstrate that WRN associates with BLM via the DHBN domain independently of the hydrophobic homodimerization patch, and that this interaction is attenuated upon mitotic arrest.

**Figure 1.**
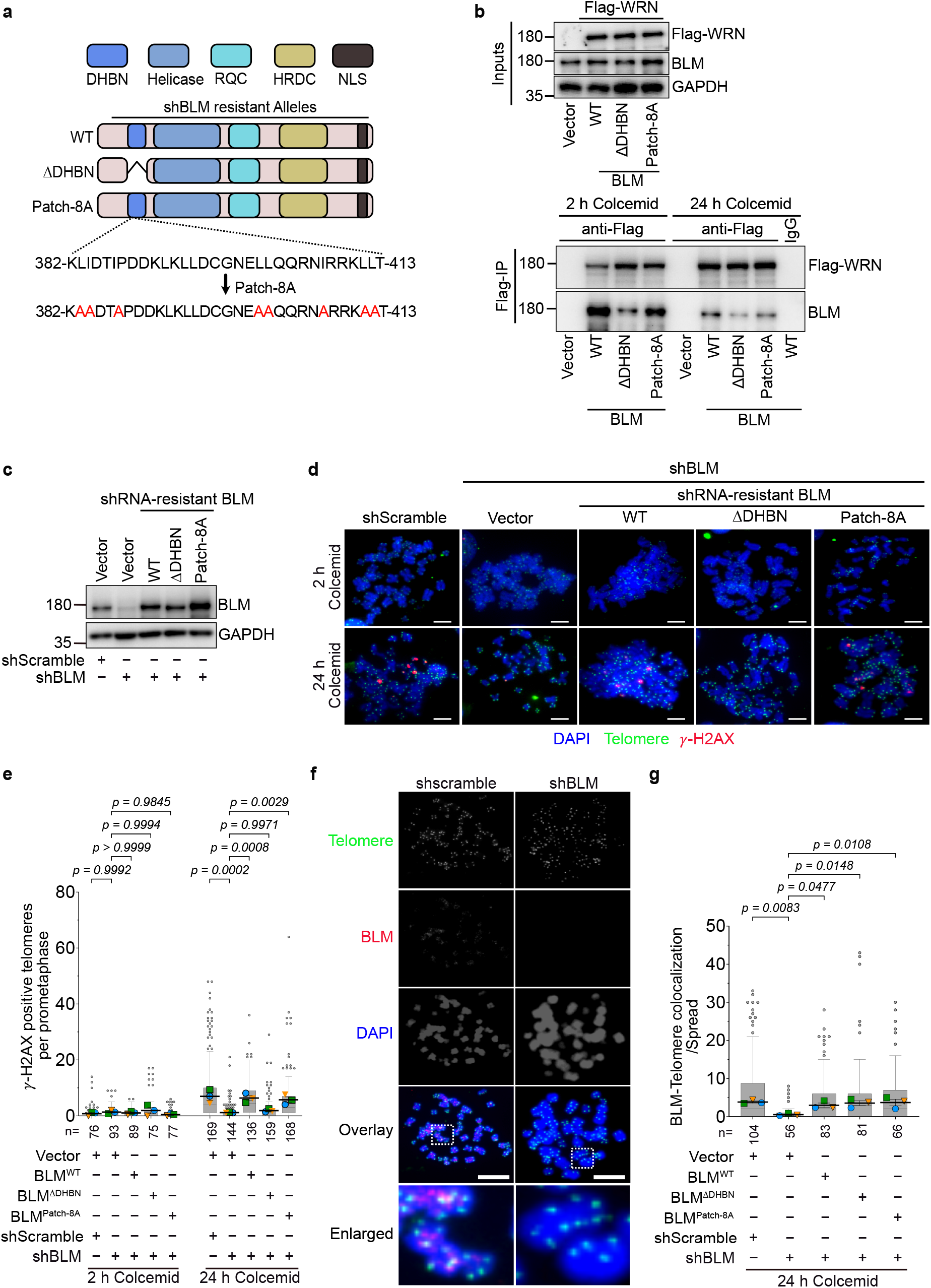
The BLM DHBN domain is essential for BLM-driven MAD telomere deprotection. **a)** Schematic of Myc-BLM constructs: wild-type (WT), a DHBN deletion mutant (ΔDHBN), and a Patch-8A mutant in which a cluster of hydrophobic residues within the DHBN domain was replaced with alanines. **b)** Co-immunoprecipitation of Myc-BLM variants with Flag-WRN in IMR-90 E6E7 hTERT cells treated with 100 ng mL^-1^ colcemid for 2 h or 24 h. Flag-WRN was immunoprecipitated with anti-Flag antibody, and co-precipitated Myc-BLM was detected using an anti-BLM antibody. **c)** Immunoblot confirming expression of Myc-BLM^WT^, Myc-BLM^ΔDHBN^, and Myc-BLM^Patch-8A^ following depletion of endogenous BLM by shBLM. **d)** Representative images of metaphase telomere deprotection-induced foci (meta-TIF) in metaphase spreads, visualized by combined γ-H2AX immunofluorescence (red), telomere FISH (green), and DAPI staining (blue). IMR-90 E6E7 hTERT cells expressing the indicated BLM variants and shBLM were treated with 100 ng mL^-1^ colcemid for 2 h or 24 h. Scale bar, 10 µm. **e)** Quantification of meta-TIF per metaphase spread corresponding to (d). The total number of metaphase spreads analyzed (n) from 3 biological replicates is indicated below each dataset. Data are presented as Tukey box plots of all individual data points, with the mean of each biological replicate superimposed as a distinct colored symbol (circles, squares, and triangles), and the mean ± s.d. of the 3 replicate means is indicated by a horizontal line with error bars. Exact p values are indicated above each comparison; one-way ANOVA followed by Tukey’s multiple-comparisons test. **f)** Representative images of BLM–telomere colocalization on mitotic chromosomes IMR-90 E6E7 hTERT cells expressing shscramble or shBLM, visualized by combined BLM immunofluorescence (red), telomere FISH (green), and DAPI (blue). Cells were treated with 100 ng mL^-1^ colcemid for 24 h before preparation of metaphase spreads. Scale bar, 10 µm. **g)** Quantification of BLM signals colocalizing with telomere signals on mitotic chromosomes in the cell lines described in (c), treated as in (f). Data are presented as in (e).

We next asked whether the DHBN domain is required for BLM-mediated MAD telomere deprotection. To address this, IMR-90 hTERT E6E7 cells were transduced with lentivirus expressing shRNA-resistant BLM constructs, followed by transduction with lentivirus encoding *BLM*-targeting shRNA (shBLM) (Figure 1c). Cells were treated with colcemid for 24 hours to induce mitotic arrest, and MAD telomere deprotection was quantified by scoring metaphase TIF (meta-TIF), defined as the colocalization of the DNA damage marker γ-H2AX with telomeric signals.

Meta-TIF levels were significantly elevated in shScramble control cells following 24 hours of colcemid treatment, confirming robust MAD telomere deprotection (Figure 1d, e). This induction was nearly completely abrogated in shBLM-expressing cells, consistent with previous reports [16] (Figure 1d, e). Notably, re-expression of BLM^ΔDHBN^, but not BLM^WT^, failed to restore meta-TIF levels, demonstrating that the DHBN domain is essential for BLM’s role in promoting MAD telomere deprotection (Figure 1d, e). Interestingly, meta-TIF levels in cells expressing BLM^Patch-8A^ were reduced relative to those in BLM^WT^-expressing cells, yet remained significantly higher than in vector control cells, indicating that BLM^Patch-8A^ retained residual MAD telomere deprotection-promoting activity (Figure 1d, e). None of these BLM variants affected meta-TIF levels under the 2-hour colcemid condition, indicating that BLM is not required to maintain protective telomere structure in interphase cells (Figure 1d, e). Collectively, these data establish that the DHBN domain is required for BLM to promote MAD telomere deprotection, and that the BLM homodimerization surface contributes to, but is not essential for, this function.

Given that DHBN deletion markedly reduced MAD telomere deprotection while BLM^Patch-8A^ retained MAD telomere deprotection-promoting activity, we investigated whether the DHBN domain contributes to telomeric targeting of BLM during prolonged mitotic arrest. We performed immunofluorescence and fluorescence in situ hybridization (IF-FISH) to quantify BLM localization at telomeres following 24 hours of colcemid treatment. BLM^WT^, BLM^ΔDHBN^, and BLM^Patch-8A^ were all detected at telomeres to a comparable extent, suggesting that the DHBN domain is dispensable for telomeric recruitment of BLM during mitotic arrest (Figure 1f, g).

### The BLM^Patch-8A^ is resistant to WRN-mediated suppression of MAD telomere deprotection

The residual MAD telomere deprotection-promoting activity of BLM^Patch-8A^ prompted us to ask whether WRN could suppress BLM^Patch-8A^-dependent MAD telomere deprotection. We co-expressed BLM^WT^ or BLM^Patch-8A^ with either empty vector, full-length WRN (WRN^WT^), or the WRN CCR (WRN^168-333^) that is sufficient for suppression of MAD telomere deprotection [15]. All constructs were expressed in cells expressing shBLM, and cells were subjected to 24-hour colcemid treatment (Figure 2a, b). As expected, overexpression of either WRN^WT^ or WRN^168-333^ in BLM^WT^-expressing cells robustly suppressed meta-TIF levels (Figure 2b). In contrast, overexpression of the same WRN constructs in BLM^Patch-8A^-expressing cells failed to affect meta-TIF levels (Figure 2b). The insensitivity of BLM^Patch-8A^-expressing cells to WRN overexpression indicates that WRN restrains BLM activity through the DHBN domain. WRN overexpression did not affect the telomeric localization of BLM variants during mitotic arrest (Figure 2c). Collectively, these findings suggest that WRN suppresses MAD telomere deprotection through a direct, yet hydrophobic patch-independent, interaction with the BLM DHBN domain, acting downstream of BLM telomere localization. These results establish the DHBN domain as a central regulatory hub for MAD telomere deprotection.

**Figure 2.**
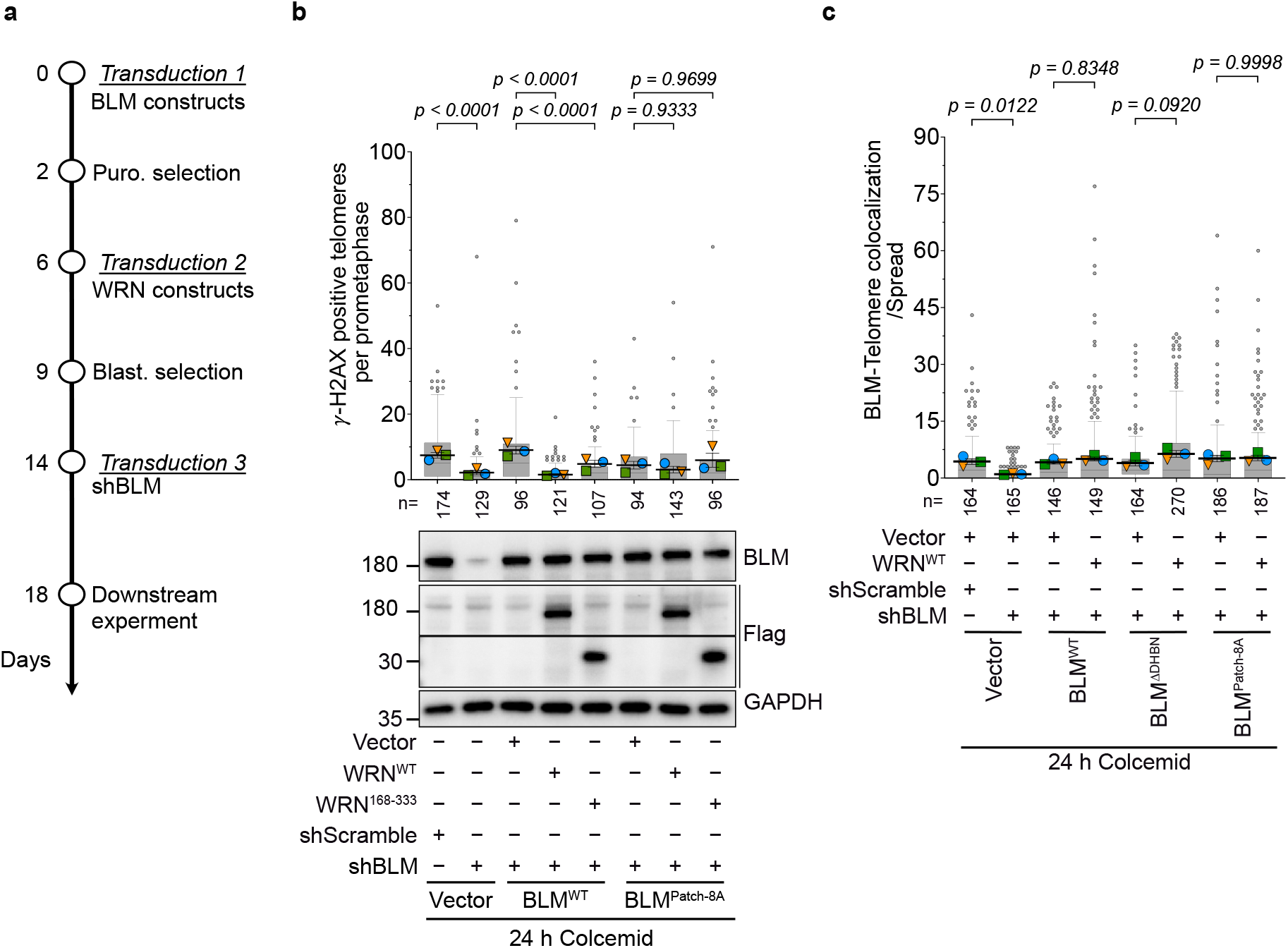
WRN-dependent suppression of MAD telomere deprotection requires an intact BLM DHBN domain. **a)** Schematic of the experimental timeline for (b). IMR-90 E6E7 hTERT cells were transduced with BLM constructs and selected with puromycin (Puro). These cells were subsequently transduced with Flag-WRN variants, selected with Blasticidin S, and finally transduced with a lentivirus encoding shBLM. **b)** Top: quantification of meta-TIF in indicated IMR-90 E6E7 hTERT cells following 24 h treatment with 100 ng mL^-1^ colcemid. Data are presented as in Figure 1e. Myc-BLM^WT^ or Myc-BLM^Patch-8A^ was co-expressed with empty vector, 4×Flag-WRN^WT^ or 4×Flag-WRN^168-333^. Bottom: representative immunoblots of whole-cell extracts from the untreated cells. **c)** Quantification of BLM signals colocalizing with telomere signals in IMR-90 E6E7 hTERT cells expressing the indicated BLM variants together with empty vector or 4×Flag-WRN^WT^. Data are presented as in Figure 1e.

### WRN-mediated suppression of BLM is specific to mitotic arrest

We next asked whether WRN’s inhibitory effect on BLM extends to other well-characterized BLM functions outside of mitotic arrest, namely the suppression of sister chromatid exchange (SCE) [19]. BLM knockdown increased SCE frequency as expected, and this phenotype was fully rescued by re-expression of BLM^WT^ (Figure 3a-c). Both BLM^ΔDHBN^ and BLM^Patch-8A^, which exhibited no dominant-negative effect on endogenous BLM (Supplementary Figure 2a-c), only partially rescued this phenotype (Figure 3a-c), consistent with the previously reported requirement for the DHBN domain in SCE suppression [20]. However, overexpression of a panel of NLS-tagged WRN constructs—including WRN^WT^, exonuclease-dead WRN (WRN^E84A^), helicase-dead WRN (WRN^K577M^), and WRN^168-333^—did not affect SCE frequency in BLM-proficient cells (Figure 3d, f). Nuclear localization of all WRN constructs was confirmed by immunofluorescence staining (Supplementary Figure 3a, b). Taken together, these results indicate that WRN-mediated inhibition of BLM is selectively operative during mitotic arrest and does not impinge upon BLM’s canonical anti-SCE function in interphase.

**Figure 3.**
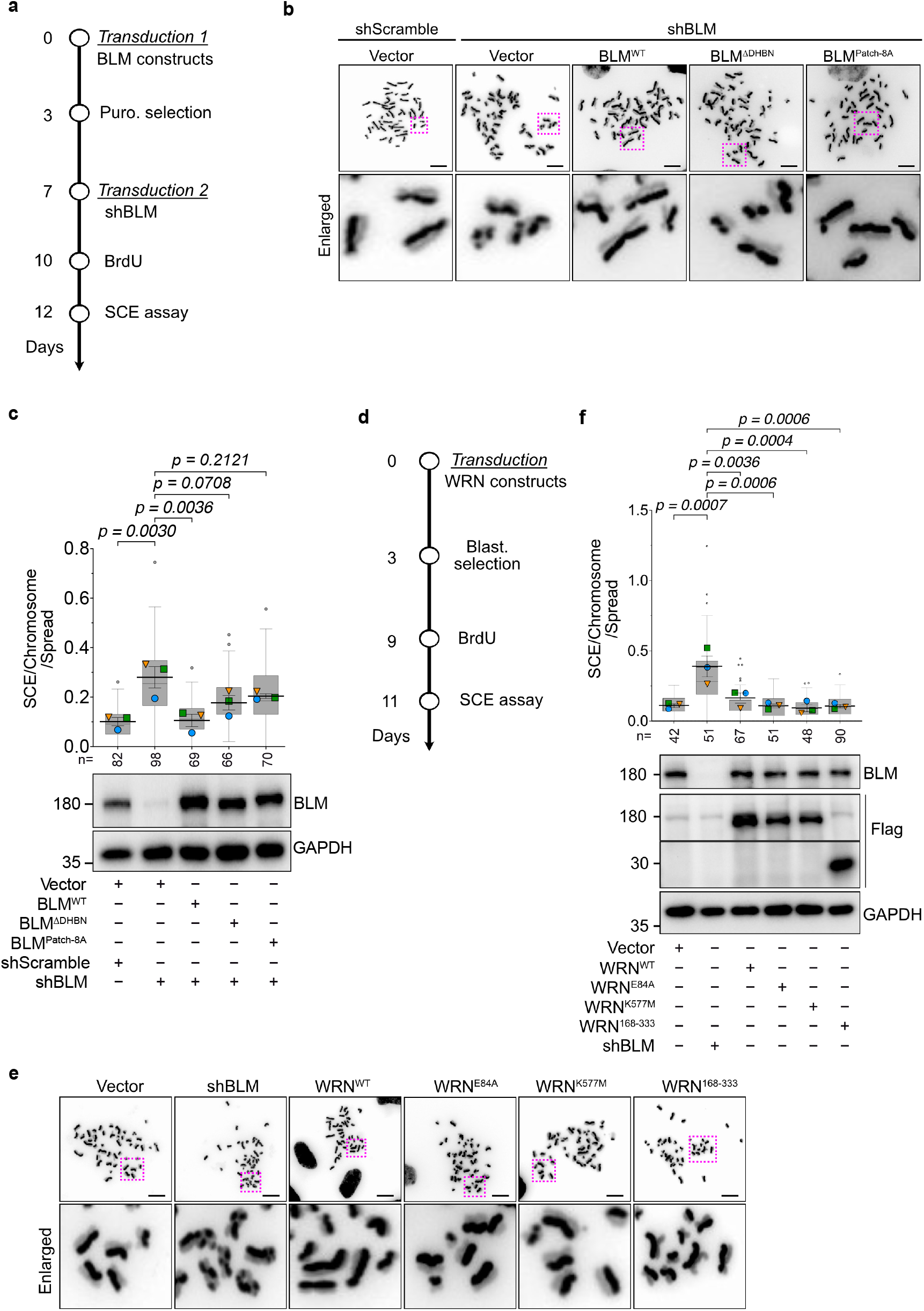
WRN does not affect the anti-SCE activity of BLM. **a)** Schematic of the experimental timeline for the sister chromatid exchange (SCE) assay. **b)** Representative images of SCE events in IMR-90 E6E7 hTERT cells expressing shBLM and the indicated BLM variants at 12 days post-BLM variant transduction. Cells were treated with BrdU for 48 h, then fixed and stained with DAPI to visualize SCE events on metaphase spreads. Dashed magenta boxes indicate the regions shown in the enlarged panels. Scale bar, 10 µm. **c)** Top: quantification of SCE events corresponding to (b). Data are presented as in Figure 1e. Bottom: immunoblot showing the knockdown efficiency of endogenous BLM and the expression levels of the indicated BLM variants. **d)** Schematic of the experimental timeline for the SCE assay in (e) and (f). **e)** Representative images of SCE events in IMR-90 E6E7 hTERT cells expressing shBLM or the indicated WRN variants. Dashed magenta boxes indicate the regions shown in the enlarged panels. Scale bar, 10 µm. **f)** Top: quantification of SCE events corresponding to (e). Data are presented as in Figure 1e. Bottom: immunoblot experiment showing the knockdown of endogenous BLM and the expression levels of the indicated WRN variants.

### GEN1 and MUS81 catalytic activities contribute to MAD telomere deprotection

Our results support a model in which the telomeric T-loop junction is processed in a manner analogous to homologous recombination intermediates during prolonged mitotic arrest. Building on this framework, we next investigated whether additional recombination-associated enzymes contribute to MAD telomere deprotection. We focused on MUS81 and GEN1, two structure-selective endonucleases with well-established roles in processing branched DNA intermediates in eukaryotic cells [25, 26]. GEN1 has been reported to resolve Holliday junction-like intermediates during anaphase [27], whereas MUS81 resolves aberrant DNA structures that persist into mitosis [28]. We constructed two independent shRNAs targeting each of *MUS81* and *GEN1* and assessed knockdown efficiency by RT-qPCR (Supplementary Figure 4a-c). As shMUS81-2 showed slightly lower knockdown efficiency and shGEN1-2 exhibited cytotoxicity and negligible knockdown efficiency, shMUS81-1 and shGEN1-1 were selected for subsequent analyses.

MUS81 depletion had no detectable effect on meta-TIF levels in the absence of prolonged mitotic arrest (Figure 4a–d: 2 h colcemid). In contrast, MUS81 depletion resulted in a reduction in meta-TIF levels relative to shScramble control cells upon mitotic arrest (Figure 4a–d: 24 h colcemid). Re-expression of shRNA-resistant MUS81^WT^, but not a catalytically inactive MUS81^D338A/D339A^ mutant [29], in shMUS81-expressing cells restored meta-TIF levels (Figure 4a–d); while both MUS81^WT^ and MUS81^D338A/D339A^ were detected at telomeres during mitotic arrest (Figure 4e, f). These results suggest that MUS81 enzymatic activity contributes to MAD telomere deprotection.

**Figure 4.**
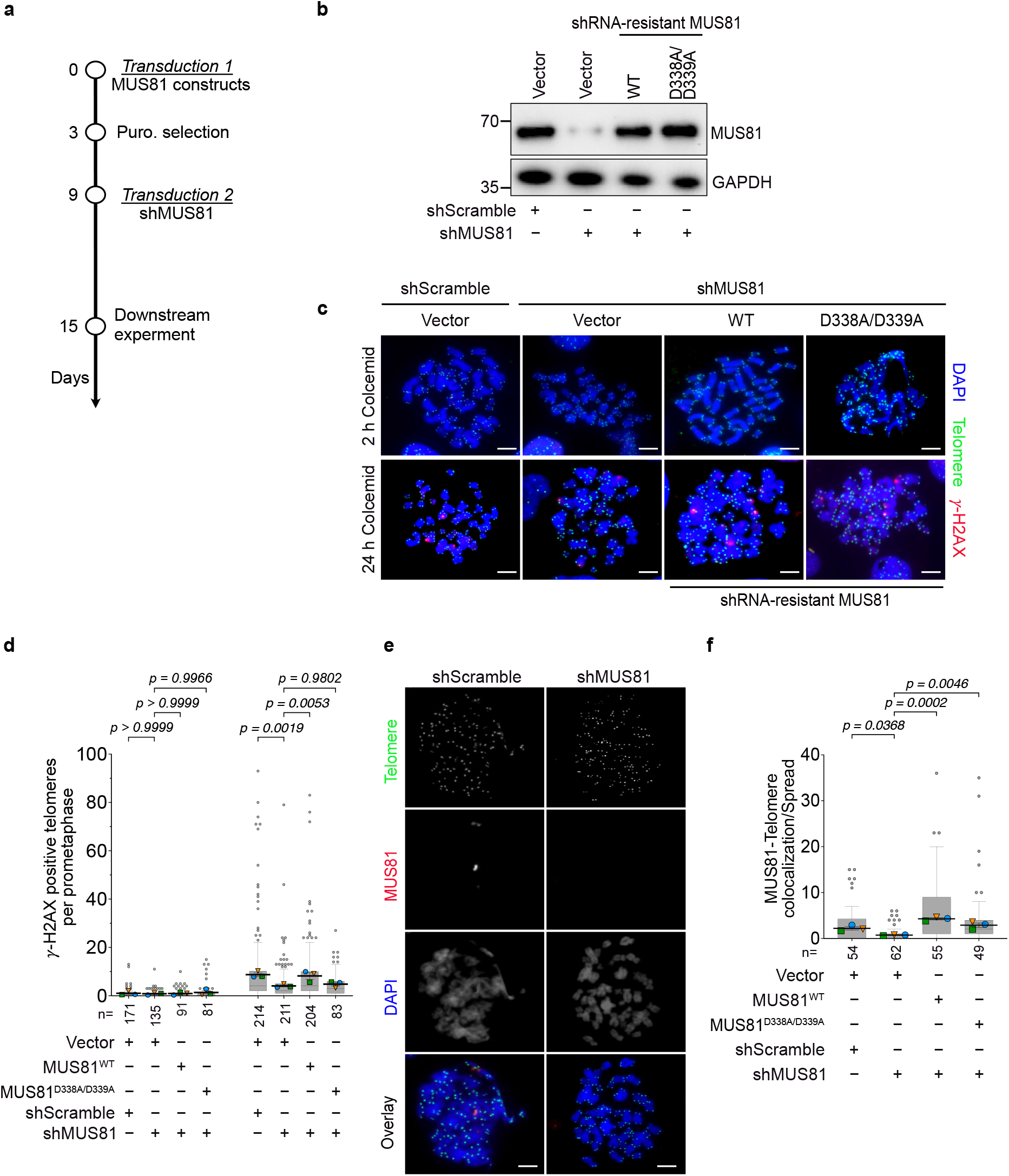
MUS81 catalytic activity contributes to MAD telomere deprotection. **a)** Schematic of the experimental timeline for (b)–(f). (**b**) Representative immunoblot analysis of whole-cell extracts from IMR-90 E6E7 hTERT cells expressing the indicated MUS81 variants and shMUS81-1, collected at 5 days post-shRNA transduction. **c)** Representative meta-TIF images of IMR-90 E6E7 hTERT cells described in (b) following 2 h or 24 h colcemid treatment. Scale bar, 10 µm. **d)** Quantification of meta-TIF on mitotic chromosome spreads corresponding to (c). Data are presented as in Figure 1e. **e)** Representative images of MUS81–telomere colocalization on mitotic chromosomes in IMR-90 E6E7 hTERT cells expressing shScramble or shMUS81-1, visualized by combined MUS81 immunofluorescence (red), telomere FISH (green), and DAPI (blue). Scale bar, 10 µm. **f)** Quantification of MUS81 signals colocalizing with telomeres on mitotic chromosomes in IMR-90 E6E7 hTERT cells expressing the indicated MUS81 variants and shMUS81-1. Data are presented as in Figure 1e.

We further explored the function of GEN1 and found that GEN1 depletion produced a pronounced reduction in meta-TIF levels upon mitotic arrest, while having a negligible effect on meta-TIF under the 2-hour colcemid control condition (Figure 5a–d). Re-expression of GEN1^WT^, but not the catalytically inactive GEN1^E134A/E136A^ mutant [26], rescued this phenotype (Figure 5a–d). Like MUS81, GEN1 variants were detected at mitotic telomeres upon prolonged mitotic arrest (Figure 5e, f). These results demonstrate that GEN1 catalytic activity also contributes to MAD telomere deprotection.

**Figure 5.**
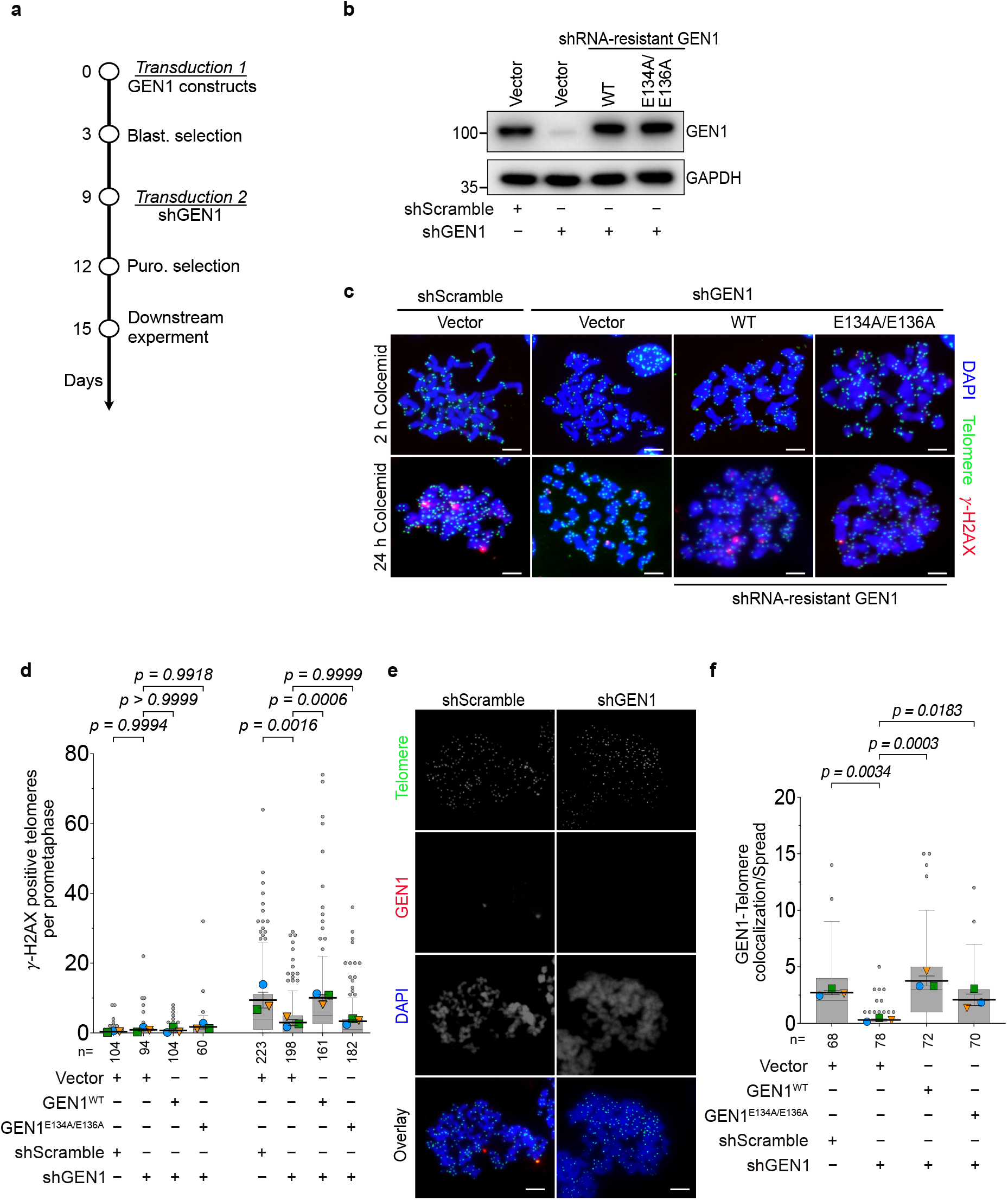
GEN1 catalytic activity promotes MAD telomere deprotection. **a)** Schematic of the experimental timeline for (b)–(f). **b)** Representative immunoblot analysis of whole-cell extracts from IMR-90 E6E7 hTERT cells expressing the indicated GEN1 variants and shGEN1-1, collected at 5 days post-shRNA transduction. **c)** Representative images of meta-TIF assay in IMR-90 E6E7 hTERT cells corresponding to (b) following 2 h or 24 h colcemid treatment. Scale bar, 10 µm. **d)** Quantification of meta-TIF in mitotic chromosome spreads corresponding to (c). Data are presented as in Figure 1e. **e)** Representative images of GEN1–telomere colocalization on mitotic chromosomes in IMR-90 E6E7 hTERT cells expressing shScramble or shGEN1-1, visualized by combined GEN1 immunofluorescence (red), telomere FISH (green), and DAPI (blue). Scale bar, 10 µm. **f)** Quantification of GEN1 signals colocalizing with telomeres on mitotic chromosomes in IMR-90 E6E7 hTERT cells expressing the indicated GEN1 variants and shGEN1-1. Data are presented as in Figure 1e.

Early studies reported that co-depletion of MUS81 or GEN1 with SLX4 prolonged mitotic duration [25], raising the possibility that depletion of these resolvases might indirectly affect the extent of MAD telomere deprotection by altering the duration of mitotic arrest. To address this potential confound, we analyzed mitotic duration in GEN1-or MUS81-depleted cells by live-cell imaging. Mitotic duration was not significantly altered by depletion of either enzyme under either unperturbed conditions or colcemid treatment (Supplementary Figure 4d, e). Collectively, these results establish that the catalytic activities of MUS81 and GEN1 contribute to the resolution of telomeric protective structures during mitotic arrest.

### TRF2 suppresses BLM-, MUS81-, and GEN1-mediated MAD telomere deprotection

We have previously shown that TRF2 protects mitotic telomeres from BLM-dependent dissolution [16]. The basic domain of TRF2 has previously been shown to interact with core histones to reinforce T-loop stability [30]. To determine whether the protective interaction between the TRF2 basic domain and core histones is attenuated during mitotic arrest, we performed a Co-IP experiment in IMR-90 E6E7 hTERT cells expressing Flag-TRF2 variants. We detected robust histone H2B and H3 signals in Flag-TRF2^WT^ immunoprecipitates (Figure 6a, b). Consistent with the previous report [30], this interaction was significantly attenuated in Flag-TRF2^ΔBD^ immunoprecipitates (Figure 6a, b). Notably, this interaction was nearly abolished upon mitotic arrest even in Flag-TRF2^WT^-expressing cells (Figure 6b and Supplementary Figure 5a, b). We further found that the interaction was abolished by pre-treatment of lysates with Benzonase (Figure 6b and Supplementary Figure 5a, b), suggesting that nucleic acids, such as telomeric DNA and RNA, stabilize it. Taken together, these data indicate that the interaction between the TRF2 basic domain and core histones is abrogated during prolonged mitotic arrest.

**Figure 6.**
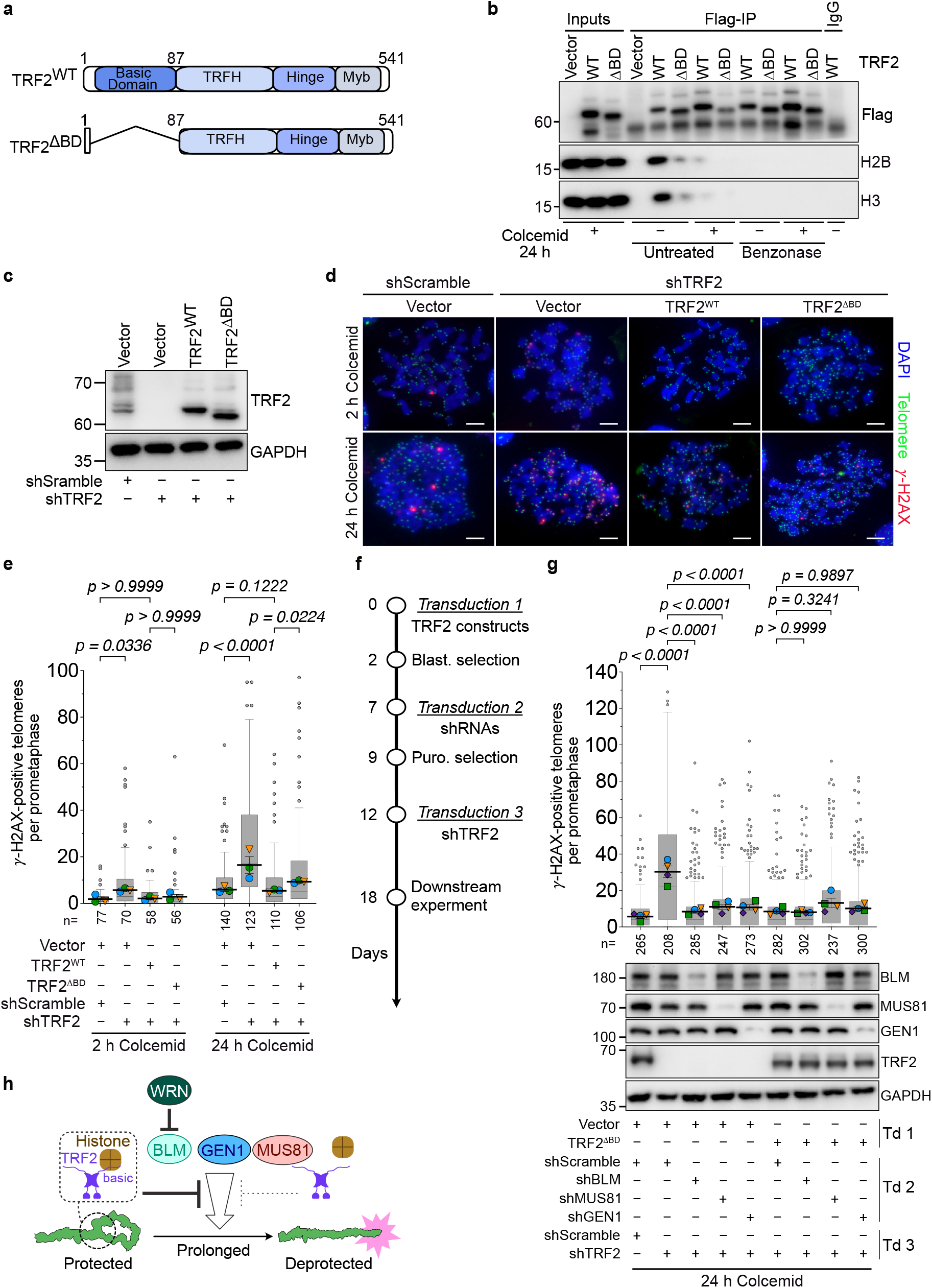
The TRF2 basic domain interacts with core histones and contributes to mitotic telomere protection. **a)** Schematic of TRF2 constructs used in this study, including wild-type TRF2 (TRF2^WT^) and the basic domain deletion mutant (TRF2^ΔBD^). TRF2^ΔBD^ lacks residues A2–G86 of human TRF2 (XP_005256180.1). **b)** Immunoblots of anti-Flag immunoprecipitates from IMR-90 E6E7 hTERT cells expressing the indicated Flag-TRF2 variants. Cells were left untreated or treated with colcemid for 24 h, and collected at 6 days post-transduction. Whole-cell lysates were incubated with or without Benzonase for 1 h at 37℃ before immunoprecipitation. **c)** Immunoblot analysis of whole-cell extracts from IMR-90 E6E7 hTERT cells expressing the indicated shRNAs and TRF2 variants, collected at 5 days post-shRNA transduction. **d)** Representative meta-TIF images of mitotic chromosome spreads from IMR-90 E6E7 hTERT cells described in (c) following 2 h or 24 h colcemid treatment. Scale bar, 10 µm. **e)** Quantification of meta-TIF corresponding to (d). Data are presented as in Figure 1e. **f)** Schematic of the experimental timeline for (g). IMR-90 E6E7 hTERT cells were sequentially transduced with empty vector or TRF2^ΔBD^, followed by shScramble, shBLM, shMUS81-1, or shGEN1-1, and finally shScramble or shTRF2-F. **g)** Top: quantification of meta-TIF in the cell lines established in (f). Data are presented as in Figure 1e (N = 4 biological replicates). Bottom: representative immunoblots of whole-cell extracts from the untreated cells. Td; transduction. **h)** Model for T-loop processing during prolonged mitotic arrest. Prolonged mitotic arrest attenuates the protective functions of TRF2, including the interaction between the TRF2 basic domain and core histones, thereby potentially rendering the T-loop junction accessible to the BTRR complex. BLM may then promote branch migration and generate Holliday junction intermediates. These intermediates are subsequently processed by either dissolution via the BTRR complex or nucleolytic resolution by the structure-selective endonucleases MUS81 and GEN1.

This result prompted us to investigate the role of the TRF2 basic domain in the suppression of BLM, MUS81, and GEN1 during mitotic arrest. IMR-90 E6E7 hTERT cells expressing shTRF2 exhibited a significant level of meta-TIF under the 2-hour colcemid condition (Figure 6c–e). These are interphase-derived TIFs carried over into mitosis [17]. Twenty-four hours of colcemid treatment further exacerbated meta-TIF, indicating robust MAD telomere deprotection in TRF2-depleted cells (Figure 6c–e). Re-expression of TRF2^WT^ or TRF2^ΔBD^ suppressed meta-TIF levels under the 2-hour colcemid condition, consistent with previous reports demonstrating that the basic domain is dispensable for the prevention of interphase TIF [1, 31] (Figure 6c–e). However, cells expressing TRF2^ΔBD^ failed to reduce the meta-TIF levels to those of TRF2^WT^ under 24-hour colcemid conditions, indicating a protective function of the basic domain specifically during mitotic arrest. We then transduced cells expressing shTRF2 and TRF2^ΔBD^ with shBLM, shMUS81-1, or shGEN1-1 to assess the contributions of these enzymes to MAD telomere deprotection in the absence of functional TRF2 (Figure 6f). In cells expressing shTRF2 without TRF2 rescue, depletion of each enzyme markedly reduced meta-TIF levels (Figure 6g), indicating that TRF2 protects mitotic telomeres from these enzymatic activities. In TRF2^ΔBD^-expressing cells, however, depletion of none of these enzymes significantly affected meta-TIF levels (Figure 6g). These results demonstrate that TRF2 protects T-loop junctions from BLM-, MUS81-, and GEN1-dependent enzymatic activities during mitotic arrest (Figure 6h), with the basic domain potentially contributing to this protection but not being solely responsible for it.

## Discussion

Our work builds upon the conceptual framework that MAD telomere deprotection involves enzymatic activities classically associated with HR and exploits the structural similarity between telomeric T-loops and HR joint molecules. We identified a physical interaction between WRN and BLM that depends on the BLM DHBN domain, positioning this domain as a potential regulatory hub. Importantly, the DHBN domain has previously been implicated in the dissolution of dHJs [20], providing a conceptual bridge between HR intermediate processing and T-loop dismantling. We also provided evidence that the HR resolvases MUS81 and GEN1 contribute to MAD telomere deprotection, further supporting the notion that telomere protective structures are processed as HR intermediates during mitotic arrest. These enzymatic activities at T-loops are stringently repressed during the normal cell cycle by TRF2 through multiple mechanisms. Our data suggest that weakening of the TRF2–histone association during mitotic arrest contributes to a permissive state for dissolution and resolution of protective T-loop junctions, although the observation that TRF2^ΔBD^ only partially phenocopies TRF2 depletion indicates that additional TRF2 domains act in parallel to maintain telomere protection during mitotic arrest.

### The DHBN domain mediates a telomere-specific BLM activity that is selectively targeted by WRN during mitotic arrest

The functional consequences of disrupting the BLM DHBN domain reveal a mechanistic distinction between BLM’s roles at mitotic telomeres and in the interphase genome. DHBN deletion nearly abolishes MAD telomere deprotection, while the BLM^Patch-8A^ mutant retains residual activity (Figure 1c-e). In contrast, in sister chromatid exchange (SCE) assays performed in BLM-depleted cells, both BLM^ΔDHBN^ and BLM^Patch-8A^ partially restore SCE suppression to a similar extent (Figure 3b, c). These results suggest that the DHBN domain harbors an additional, mitotic telomere-specific function that is maintained in the BLM^Patch-8A^ mutant. Because both BLM^ΔDHBN^ and BLM^Patch-8A^ were detected at mitotic telomeres (Figure 1f, g), this uncharacterized DHBN function is likely required for processing steps downstream of telomeric recruitment. Interestingly, WRN overexpression did not affect MAD telomere deprotection in cells expressing BLM^Patch-8A^ (Figure 2b), even though the WRN–BLM^Patch-8A^ interaction was largely preserved (Figure 1b). We therefore propose that WRN targets this telomere-specific, uncharacterized function of the DHBN domain, which becomes structurally insensitive to WRN-mediated suppression in the BLM^Patch-8A^ mutant (Supplementary Figure 6). This could explain how WRN selectively dampens telomere-restricted BLM activity, which is unmasked during mitotic arrest. Consistent with this interpretation, WRN overexpression did not affect BLM-dependent SCE suppression at non-telomeric loci (Figure 3e, f). Although the precise mechanism by which WRN specifically restrains BLM function at mitotic telomeres remains to be determined, the WRN– BLM interaction mediated through the DHBN domain emerges as a regulatory module that imposes spatial and temporal control over telomeric BLM activities.

### Protective telomere structures are processed as homologous recombination intermediates during mitotic arrest

Our analyses of MUS81 and GEN1 further extend this framework. Both enzymes contribute to MAD telomere deprotection (Figures 4 and 5). Importantly, the involvement of both the BTRR dissolvase complex [16] and the structure-selective resolvases underscores a central unresolved question: the precise nature of the telomeric substrate during MAD telomere deprotection. While all these enzymes can catalyze D-loop disruption or cleavage *in vitro* [32], it is noteworthy that BLM knockdown almost completely suppresses MAD telomere deprotection (Figure 1c-e) [16], suggesting that MUS81 and GEN1 have negligible effects on telomere protective structures in the absence of BLM activity. Similarly, WRN overexpression robustly abolishes MAD telomere deprotection even in the presence of functional MUS81 and GEN1 (Figure 2b) [15]. These results suggest that BLM functions as the primary enzyme that either unwinds D-loops at T-loop junctions or generates single-and double-Holliday junction intermediates through branch migration during mitotic arrest (Supplementary Figure 7) [17]. Previous *in vitro* experiments have shown that the TOP3A–RMI1–RMI2 complex enhances BLM’s D-loop unwinding activity by regulating the directionality of BLM’s helicase activity [33], implicating the BTRR complex in this unwinding reaction (Supplementary Figure 7). Branch migration that produces Holliday junctions from single D-loops may also rely on the BTRR complex and/or the uncharacterized function of the DHBN domain. These secondary products are either dissolved by BTRR or resolved by MUS81 and GEN1 (Supplementary Figure 7).

### The TRF2 basic domain–histone interaction is attenuated during mitotic arrest, partially licensing T-loop processing by recombination enzymes

The basic domain of TRF2 has previously been shown to protect telomeres from recombination-associated enzymatic activities [31, 34, 35], potentially through its interaction with core histones [30]. Consistent with this, our results demonstrate that TRF2 suppresses the activities of BLM, MUS81, and GEN1 on telomere protective structures during mitotic arrest. We further show that the TRF2 basic domain–histone interaction is nearly abolished during prolonged mitotic arrest (Figure 6b). Sensitivity of this interaction to Benzonase treatment indicates that the TRF2–histone association is stabilized by telomeric chromatin context, including telomeric DNA and RNA, rather than constituting a stable protein–protein interaction. Together with the recent finding that phospho-mimetic mutations within the TRF2 basic domain near the histone-binding motif strongly promote MAD telomere deprotection [16], these observations suggest that mitotic phosphorylation within the basic domain loosens TRF2–histone contacts, thereby contributing to a permissive state for enzymatic processing of T-loop junctions (Figure 6h). This provides a plausible upstream mechanism by which telomeres become accessible to recombination enzymes during mitotic arrest without invoking loss of TRF2 binding to telomeric DNA.

However, the observation that TRF2^ΔBD^ only partially phenocopies TRF2 depletion, and that depletion of BLM, MUS81, or GEN1 did not significantly affect meta-TIF levels in TRF2^ΔBD^-expressing cells, suggests that the basic domain is not the sole determinant of TRF2-mediated protection during mitotic arrest. The interaction between TRF2 and RAP1, mediated through the TRF2 Hinge domain [36, 37] as well as the iDDR region of TRF2 [1], represents additional plausible candidates in this regard.

The contrasting phenotypic severity between TRF2^ΔBD^ and phospho-mimetic mutations within the basic domain [24] is noteworthy. Whereas deletion of the entire basic domain produces a relatively mild MAD telomere deprotection phenotype, phospho-mimetic substitutions at specific residues within the same domain elicit substantially stronger telomere deprotection [24]. This discrepancy raises the intriguing possibility that the basic domain harbors both suppressive and pro-deprotection activities, such that its complete deletion simultaneously abrogates both functions, thereby attenuating the net phenotypic consequence. In this model, mitotic phosphorylation would serve not merely to relieve suppressive TRF2–histone contacts but to actively promote T-loop processing, potentially by converting the basic domain from a protective to a pro-deprotection module. Resolving this possibility will require systematic dissection of the individual residues and interactions within the basic domain that contribute to each function.

Taken together, our study defines MAD telomere deprotection as a regulated, multi-enzyme process that repurposes HR-associated activities toward telomere protective structures while imposing strict contextual control through protein–protein interactions and chromatin state. By identifying the BLM–WRN and TRF2–histone associations as plausible inhibitory nodes in this pathway, we provide a mechanistic framework explaining how telomeres are selectively deprotected during mitotic arrest without compromising genome-wide stability. More broadly, these findings demonstrate how cells can reprogram recombination enzymes to remodel specialized DNA structures in a cell cycle-specific manner, highlighting telomeres as dynamic, context-dependent platforms for DDR activation rather than passive and static protective structures.

### Limitations of the study

While our findings provide significant mechanistic insight into MAD telomere deprotection, several important limitations warrant consideration. First, our study relies primarily on quantifying meta-TIF levels as a surrogate marker of telomere deprotection. Although this assay is widely used and provides a robust readout of telomere-associated DNA damage signaling, it remains an indirect measure of T-loop disruption. Future studies employing biochemical reconstitution systems or high-resolution structural imaging approaches will be required to directly assess the structural dynamics of telomeric DNA and confirm T-loop disruption events. Second, the experimental framework is based on prolonged mitotic arrest induced by microtubule depolymerization. While this approach enables efficient induction of MAD telomere deprotection and facilitates mechanistic interrogation, it may not fully recapitulate the conditions of physiological mitotic delay or arrest. Third, the interpretation of the TRF2^ΔBD^ rescue experiments is constrained by the requirement for three sequential lentiviral transductions, which may have introduced technical variability that limits the sensitivity of this experimental system. Furthermore, more systematic mutational approaches targeting individual residues within TRF2 will be required to fully dissect the contribution of its functional domains to the regulation of MAD telomere deprotection.

## Materials and methods

### Cell lines

IMR-90 normal diploid human fibroblast cells (I90-26, Coriell Institute) were transduced with retroviruses encoding the HPV16 E6 and E7 oncoproteins (pLXSN3-16E6E7) and human telomerase reverse transcriptase (pWZL-hTERT) to obtain IMR-90 E6E7 hTERT cells [15]. Cells were cultured in Dulbecco’s Modified Eagle Medium (DMEM) supplemented with 10% fetal bovine serum (FBS), 7.5% NaHCO_3_, 100 U mL^-1^ penicillin-streptomycin, 200 mM L-glutamine, and 5 μg mL^-1^ Plasmocin (InvivoGen) and maintained at 37°C in 5% CO_2_ and 3% O_2_. Mitotic arrest was induced by treatment with 100 ng mL^-1^ KaryoMAX Colcemid (15212012, Thermo Fisher Scientific).

### Plasmid construction

All truncations and point mutations in the protein of interest were first modeled in SnapGene software (version 5.0.8) and then introduced by site-directed PCR mutagenesis, followed by confirmation by Sanger sequencing. Briefly, complementary DNAs (cDNAs) encoding human MUS81 (NP_079404.3) and GEN1 (NP_054197745.1) were obtained by reverse transcription of IMR-90 total RNA and cloned in-frame into third-generation lentiviral plasmid vectors, either downstream of a blasticidin S resistance gene and self-cleaving T2A peptide sequences, or upstream of an internal ribosome entry site (IRES) sequence and a puromycin resistance gene. Short hairpin RNAs (shRNAs) targeting *MUS81* and *GEN1* were cloned into the pLKO.1 vector by ligation of annealed oligonucleotides. The shRNA target sequences used in this study were as follows: shScramble, 5‘-CTAAGGTTAAGTCGCCCTCG-3‘; shTRF2, 5‘-GCGCATGACAATAAGCAGATT-3‘ (TRCN0000004811) [11]; shBLM, 5‘-TGCCAATGACCAGGCGATC-3‘ [16]; shMUS81-1, 5‘-ACACTGCTGAGCACCATTAAG-3‘ [38]; shMUS81-2, 5‘-GCAGGAGCCATCAAGAAT-3‘; shGEN1-1, 5‘-CCAGATGAAGTAATGAGCTTT-3‘ (TRCN0000051878); and shGEN1-2, 5‘-CAATACTTCTGTCCCTTATTC-3‘ (TRCN0000425804). shRNA-resistant silent mutations were introduced as follows: TRF2, 5’-G aGa ATG ACt ATc tct cGc cT-3’; BLM, 5‘-c GCt AAc GAt CAa GCc ATt-3′ [16]; MUS81, 5‘-ACt CTt CTt tct ACt ATc AAa-3‘; and GEN1, 5‘-CCt GAc GAg GTt ATG tct TTc-3‘, where spaces separate codons and lowercases denote silent mutations. The BLM (NP_000048.1) mutations were as follows: ΔDHBN, deletion of residues D362–V414; and Patch-8A, L382A/I383A/I386A/L400A/L401A/I406A/L410A/L411A [20]. All plasmids used in this study are listed in Table 1.

**Table 1.** List of Plasmids.

| **Plasmid number (pMTH)** | **Name** | **Source** |
| --- | --- | --- |
| 127 | pLKO.1-shScramble | Addgene # 1864 |
| 285 | pLKO.1-shTRF2-F (TRCN0000004811) | [11] |
| 745 | pLKO.1-shBLM-3 | [16] |
| 1481 | pLenti-IRES-bla-gsgT2A | This study |
| 1660 | pLenti_NLS_4Flag_WRN(168-333)WT_IRES_blast | This study |
| 1743 | pLenti-1xMyc-BLM_shR-IRES(short)-puro | This study |
| 1748 | pLenti-1xMyc-BLMΔDHBN-IRES(short)-puro | This study |
| 1760 | pLenti-1xMyc-BLM(Patch-8A)-IRES-puro | This study |
| 1788 | pLenti-IRES-Puro-Vec | This study |
| 1794 | pLenti-Blast-gsgT2A-4xFL-Full length WRN | [15] |
| 1947 | pLKO.1-shGEN1-1(TRCN0000051878) | This study |
| 1948 | pLKO.1-shGEN1-2 (TRCN0000425804) | This study |
| 1949 | pLKO.1-shMUS81-1 | [38] |
| 1950 | pLKO.1-shMUS81-2 | This study |
| 1960 | pLent-bla-gsgT2A-TRF2 WT shResistant | This study |
| 1961 | pLent-bla-gsgT2A-TRF2 ΔBD shResistant | This study |
| 2067 | pLenti-bla-gsgT2A-hWRN-E84A_RshRNA | This study |
| 2068 | pLenti-bla-gsgT2A-hWRN-K577M_RshRNA | This study |
| 2074 | pLenti-myc-MUS81-shR1&2-IRES-Puro | This study |
| 2130 | pLenti-Blast-gsgT2A-3xFlag-TRF2_WT_shRes | This study |
| 2131 | pLenti-Blast-gsgT2A-3xFlag-TRF2 ΔBasic_shRes | This study |
| 2174 | pLenti-Bla-gsgT2A-3xFlag-GEN1_shR1 | This study |
| 2207 | pLenti-myc-MUS81-D338AD339A_shR1&2-IRES-Puro | This study |
| 2223 | pLenti-Bla-gsgT2A-3xFlag-GEN1-E134A-E136A_shR1 | This study |

### Lentiviral packaging and transduction

Packaging (12260, Addgene) and envelope (8454, Addgene) plasmids, generously provided by Didier Trono and Bob Weinberg, respectively, were used to produce lentiviral particles in Lenti-X 293T cells (632180, Takara Bio) in antibiotic-free medium. Twenty-four hours after co-transfection of lentiviral transfer, packaging, and envelope plasmids, the medium was replaced with fresh medium containing antibiotics, and viral supernatants were collected and filtered through a 0.45 μm membrane at 48 and 72 hours post-transfection. To determine the functional titer of the lentiviral stocks, a serial dilution assay was performed. IMR-90 E6E7 hTERT cells were transduced with a 3-to 5-fold serial dilutions of the viral supernatant in the presence of 8 µg mL^-1^ polybrene. Forty-eight hours post-transduction, cells were subjected to the corresponding antibiotic selection to determine the minimum viral dose required for 100% cell survival under selection. Target cells were then transduced with the minimal dose of viral supernatant supplemented with 8 µg mL^-1^ polybrene for 48 hours. For stable expression of WRN, GEN1, or TRF2 variants, transduced cells were selected with 10 µg mL^-1^ Blasticidin S (Funakoshi) for at least 5 days before subsequent procedures, including a second transduction. For stable expression of MUS81 or BLM variants, transduced cells were selected with 1 µg mL^-1^ puromycin (ChemCruz) for at least 3 days before subsequent procedures. For shRNA-mediated knockdown of BLM, MUS81, GEN1, and TRF2, cells were transduced with lentiviruses produced from pLKO.1-shBLM (pMTH745), pLKO.1-shMUS81-1 (pMTH1949), pLKO.1-shGEN1-1 (pMTH1947), or pLKO.1-shTRF2-F (pMTH285). Transduced cells were selected with 1 µg mL^-1^ puromycin (ChemCruz) beginning at 2 days post-transduction for at least 3 days. For the shGEN1-1 lentivirus preparation, the viral supernatant was concentrated using a Lenti-X Concentrator (631232, Clontech) according to the manufacturer’s instructions.

### Immunoblotting (Western blotting)

Cells were washed with ice-cold 1× PBS, collected by cell scrapers in a small volume of residual 1× PBS, and transferred to pre-chilled 1.5 ml microcentrifuge tubes. Cells were then pelleted at 40,000 rpm for 5 minutes at 4°C. Cell pellets were lysed in lysis buffer (1× Laemmli buffer supplemented with 2% β-Mercaptoethanol) at 1 x 10^6^ cells mL^-1^. Lysates were boiled at 105°C for 5 minutes and then cooled to room temperature (RT). Lysates were resolved by SDS-PAGE using 10% polyacrylamide gels prepared with the TGX™ FastCast™ Acrylamide Starter Kit (10%; 1610172, Bio-Rad). Electrophoresis was performed in SDS running buffer (25 mM Tris, 192 mM glycine, 0.1% SDS) at 100 V for 90 min. Proteins were transferred to 0.45 μm or 0.22 μm PVDF membranes (Immobilon-P, IPVH85R, Merck Millipore), pre-activated in 100% methanol (15 seconds) or 100% ethanol (1 minute), using a wet transfer system (1703930JA, Bio-Rad). Membranes were blocked in 5% skim milk in 1× TNT buffer for 1 hour at RT, except for membranes probed for BLM or Histones, which were blocked in Blocking One buffer (03953-95, Nacalai). Blocked membranes were incubated overnight at 4℃ with primary antibodies (see Table 2) with gentle agitation. The following day, membranes were washed three times in 1× TNT buffer (0.1 M Tris-HCl [pH 7.5], 0.15 M NaCl, 0.05% Tween20) for 5 minutes per wash. Membranes were then incubated with the appropriate secondary antibodies at the dilutions indicated in Table 2 for 1 hour at RT with gentle agitation. Following removal of the secondary antibody solution, membranes were washed three times in 1× TNT buffer for 5 minutes per wash. Membranes were incubated with a chemiluminescent HRP Substrate (Chemi-Lumi One Super, 02230-14, Nacalai), and chemiluminescent signals were captured using a ChemiDoc MP Imaging System (17001402JA, Bio-Rad). Images are presented as acquired, except for cropping. Raw membrane images are provided in the Supplementary Information.

**Table 2.** List of antibodies.

| **Antigen** | **Manufacturer** | **Catalog Number** | **Lot Number** | **Host Species** | **Dilution** |
| --- | --- | --- | --- | --- | --- |
| Flag | Sigma | F1804 | 0000466000 | Mouse | 1:1000 (WB)<br>1:200 (IF) |
| BLM | Novous Biological | NB100 | NB100-214 | Rabbit | 1:1000 (WB)<br>1:200 (IF) |
| TRF2 | Abcam | AB13579 | 1013591-2 | Mouse | 1:1000 |
| MUS81 | Santa Cruz | Sc-53582 | A2026 | Mouse | 1:1000 (WB)<br>1:200 (IF) |
| GEN1 | Novus Biologicals | S09-1B8 | 250713 | Rabbit | 1:1000 (WB)<br>1:200 (IF) |
| γ-H2AX | Biolegend | 613402 | B423646 | Rabbit | 1:200 |
| H3 | Selleck | G22B21 | F0057 (01) | Rabbit | 1:1000 |
| H2B | Selleck | F0784 | F078401 | Rabbit | 1:1000 |
| GAPDH | MBL | M171-3 | 009 | Mouse | 1:1000 |

### Immunoprecipitation and Co-Immunoprecipitation

IMR-90 E6E7 hTERT cells expressing 4×Flag-WRN and Myc-BLM constructs or Flag-TRF2 variants were seeded in 15 cm dishes and cultured for 24 hours, after which cells were treated with or without colcemid for an additional 24 hours. Following induction of mitotic arrest, the culture medium was removed, and cells were washed twice with ice-cold 1× PBS before being collected with a cell scraper. An anti-Flag antibody (F1804, Sigma-Aldrich) was conjugated to Dynabeads Protein G (10003D, Invitrogen) by incubating 30 μl of beads with 2 μg of anti-Flag antibody in 100 μl of an IP blocking buffer [1× PBS containing 0.5% (w/v) BSA and 0.1% (v/v) Triton X-100] overnight at 4 °C with gentle rotation. Following antibody conjugation, the beads were washed three times with IP blocking buffer. Concurrently, each cell pellet was lysed in 500 μl of lysis150 buffer [20 mM Tris-HCl, pH 7.5, 2.5 mM MgCl2, 150 mM KCl, 0.5% NP-40, 1 mM DTT, 0.2 Mm PMSF, and 1× cOmplete EDTA-free protease inhibitor cocktail (Nacalai)] for 1 hour on ice. Lysates were centrifuged at 12,000 × *g* for 30 minutes to pellet cell debris, and the supernatants were transferred to fresh tubes. Following protein quantification by the Bradford assay, 50 μg of total protein was reserved as input for each sample. For immunoprecipitation, anti-Flag antibody-conjugated Dynabeads were added to 2 mg of total protein and incubated at 4 °C overnight with gentle rotation. Following three washes in the lysis150 buffer, beads were resuspended directly in 2× SDS sample buffer (supplemented with 5% β-Mercaptoethanol, 1× PMSF, and 1× cOmplete inhibitor cocktail) and boiled at 105 °C for 5 minutes to elute bound proteins. Five microliters of each eluate and the reserved input samples were resolved by SDS-PAGE and analyzed by standard immunoblotting. For TRF2–histone interaction experiments, 2 mg of protein lysate was either left untreated or incubated with 125 U mL^-1^ Benzonase (70746, Novagen) in the presence of 2 mM MgCl_2_. Lysates were incubated at 37 °C for 1 hour and subsequently centrifuged at 16,000 × g for 20 min at 4 °C. The supernatant was subsequently incubated with anti-Flag antibody-conjugated Dynabeads, and immunoprecipitation was performed as described above.

### Immunofluorescence and Fluorescent in Situ Hybridization (FISH)

Meta-TIF analysis was performed as previously described [39], with minor modifications. IMR-90 E6E7 hTERT cells were treated with colcemid for 2 or 24 hours. Cells were trypsinized, pelleted, and resuspended in 0.2% hypotonic KCl solution for 10 minutes, then centrifuged onto Superfrost Plus glass slides (Fisher Scientific) using a Cytospin 4 cytocentrifuge (Thermo Scientific) at 1,200 rpm for 10 minutes. Chromosome spreads were fixed with 4% formaldehyde in 1× PBS for 10 minutes, permeabilized with KCM buffer [120 mM KCl, 20 mM NaCl, 10 mM Tris, pH 7.5, 0.1% Triton X-100] for 15 minutes at RT, and blocked in ABDIL buffer [150 mM NaCl, 20 mM Tris pH7.4, 0.1% Triton X-100, 2% BSA, 0.2% Fish Gelatin] containing 100 μg mL^-1^ RNase A for 15 minutes at 37 °C. Slides were incubated with an anti-γ-H2AX (pSer139) antibody (613402, Clone 2F3, Biolegend) at a 1:200 dilution in ABDIL buffer for 1 hour at 37℃, followed by three washes with 1x PBST. Slides were then incubated with an Alexa Fluor 568-conjugated anti-mouse secondary antibody (A11031, Invitrogen) at a 1:1000 dilution in ABDIL for 30 minutes at RT. After three washes with 1× PBST, slides were fixed with 4% formaldehyde in 1x PBS for 10 minutes at RT. Slides were dehydrated through an ethanol series [70%, 95%, 100% (v/v)]. Then, they were incubated with a Tel-C FAM-(CCCTTAA)3 PNA probe (F1001, Panagene) at 1.75 µg mL^-1^ for 8 minutes at 80 °C, followed by overnight hybridization at RT or for 2 hours at 37℃ in a humidified chamber. Slides were washed twice with PNA wash buffer A [70% Formamide, 10 mM Tris, pH 7.5] and three times with PNA wash buffer B [25 mM Tris, pH 7.5, 3 mM NaCl, 0.8% Tween-20]. DAPI was included in the final wash to counterstain chromosomal DNA. Slides were rinsed several times with deionized water and dehydrated through an ethanol series as described above. Slides were mounted with an in-house antifade mounting medium [4% n-propyl gallate (29303-92, Nacalai), 100 mM Tris-HCl pH 8.7, 90% glycerol (17018-25, Nacalai)]. Images of metaphase spreads were acquired using a 100× oil-immersion objective lens (PlanApo, NA 1.45) on a BZ-X710 fluorescence microscope (Keyence). Images were analyzed using the Hybrid Cell Count and Macro Cell Count software modules (BZ-X Analyzer, version 1.3.1.1, Keyence) for automated quantification. Outlier values (spreads with more than 184 or fewer than 50 telomere signals) were excluded from all analyses.

### Sister chromatid exchange (SCE) assay

The SCE assay was performed as previously described [40] with minor modifications. Briefly, cells (1 × 10⁵ per well) were seeded in 6-well plates, cultured for 24 hours, and then incubated with 10 μg mL^-1^ BrdU for 48 hours. Two hours before collection, cells were treated with 100 ng mL^-1^ colcemid to induce mitotic arrest. Cells were harvested by trypsinization, subjected to hypotonic treatment (0.075 M KCl, 10 minutes, RT), and fixed twice with ice-cold methanol: acetic acid (3:1). Chromosome spreads were prepared on humidified slides and allowed to age in the dark for at least 2 days. Slides were rehydrated in 1× PBS, stained with Hoechst 33258 (10 μg mL^-1^ in 2× SSC), UV-irradiated (365 nm, 1 hour) using UV Transilluminator (LM-26E, UVP), and incubated in preheated 2× SSC at 60 °C for 20 minutes. After DAPI staining in 2x SSC, slides were rinsed with deionized water, dehydrated through an ethanol series, mounted with in-house antifade mounting medium, and imaged at 100× magnification using a BZ-X710 fluorescence microscope (Keyence). Images were converted to 16-bit format, and SCE events were manually scored using ImageJ (v1.54g).

### RNA extraction and RT-qPCR

Total RNA was extracted at 6 days post-transduction using the RNeasy Plus Mini kit (Qiagen) according to the manufacturer’s protocol. A total of 500 ng of extracted RNA was used as template for cDNA synthesis. cDNA was synthesized using the following components: 0.2 mM dNTP (TakaRa Bio), 1 U/μL ribonuclease inhibitor (porcine liver; TaKaRa Bio), 0.25 U/μL AMV reverse transcriptase XL (TaKaRa Bio), and 0.125 μM of an oligo (dT) primer (Eurofins). Reverse transcription was performed under the following conditions: 30 °C for 10 minutes, 42 °C for 30 minutes, and 95 °C for 5 minutes. Quantitative PCR (qPCR) was subsequently performed using SYBR Green qPCR Master Mix (A66732, Thermo Fisher Scientific) with gene-specific primers. PCR amplification was performed using a StepOnePlus Real-Time PCR System (Applied Biosystems) with the following cycling conditions: initial denaturation at 95 °C for 15 seconds, followed by 40 cycles of denaturation at 95 °C for 15 seconds and annealing/extension at 60 °C for 60 seconds. Each sample was analyzed in technical duplicates. The expression level of each target gene was normalized to that of the reference gene *GAPDH*. For graphical representation, the normalized expression values were further normalized to the corresponding value in the shScramble control. The primer sequences used are listed in Table 3.

**Table 3.**
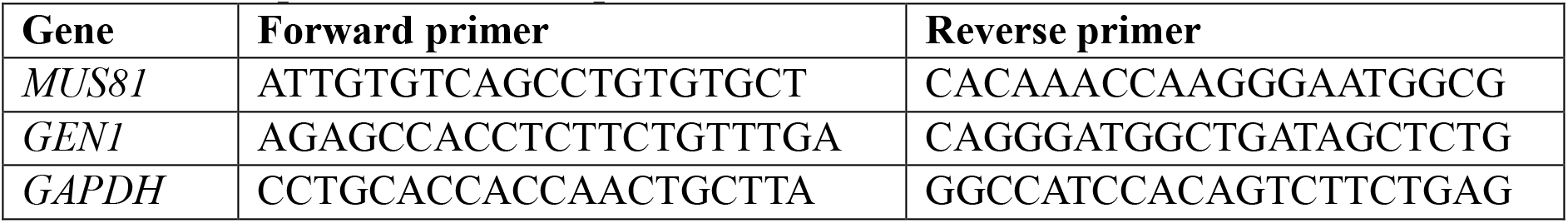
List of primers used for qPCR.

### Live cell imaging (LCI)

For live-cell imaging, 25,000 cells per well were seeded in a 12-well plate and cultured for 24 hours. Cells were treated with 100 ng mL^-1^ colcemid to induce mitotic arrest before imaging began. Time-lapse imaging was performed over 75 hours using a 10x objective lens (Plan Apo, NA 0.45) on a BZ-X710 fluorescence microscope, equipped with a stage-top incubator (INUG2-KIW, Tokai Hit) maintained at 37℃ and 5% CO_2_. Cell fate and mitotic duration were manually determined based on cell morphology, including cell rounding and nuclear envelope breakdown. Time-lapse images were compiled into video files using BZ-X Analyzer (version BZ-X_Analyzer.exe 1.3.1.1). The resulting videos were analyzed using QuickTime Player (version 10.5 (1015.4.1) and ImageJ software (v1.54g).

### Immunofluorescence

Cells were seeded on coverslips coated with Alcian Blue 8GX (A5268, Sigma-Aldrich). After 24 hours, cells were fixed with 4% paraformaldehyde in 1× PBS for 15 minutes at RT and washed 3 times with 1× PBS. Fixed cells were permeabilized for 5 minutes at RT in the dark with 0.2% Triton X-100 supplemented with 0.02% skim milk (Nacalai) and 0.02% BSA (Sigma-Aldrich) in 1× PBS. After one rinse with 1× PBS and one wash with PBST (0.1% Tween-20 in 1× PBS), cells were incubated for 45 minutes at RT with an anti-Flag primary antibody (F1804, Sigma-Aldrich) diluted 1:200 in PBST. Following three washes with PBST, cells were incubated with an Alexa Fluor 568-conjugated anti-mouse secondary antibody (A11031, Invitrogen) for 45 minutes at RT in the dark. Cells were then washed three times with PBST, and 0.1 μg mL^-1^ DAPI was added to the final wash. Cells were then rinsed with deionized water, air-dried, and mounted with in-house antifade mounting medium. Images of interphase nuclei were acquired using a 100× oil-immersion objective lens (PlanApo, NA 1.45) on a BZ-X710 fluorescence microscope (Keyence). For each condition, interphase cells were quantified using the Hybrid Cell Count and Macro Cell Count software modules (BZ-X Analyzer, version 1.3.1.1, Keyence) for automated counting.

### Statistical analysis and figure preparation

Statistical analyses were performed using GraphPad Prism (version 11.0.2 [100]). Unless otherwise indicated, all experiments were performed in three independent biological replicates. These replicate means were used for subsequent statistical analyses. Comparisons among multiple groups were performed using one-way analysis of variance (ANOVA) followed by Tukey’s multiple-comparisons test. Tukey box plots display the distribution of all individual measurements, with the center line indicating the median. The mean of each independent biological replicate is superimposed on the box plots and is denoted by blue circles (replicate 1), green squares (replicate 2), and orange triangles (replicate 3), allowing visualization of inter-experimental reproducibility. The total number of data analyzed (n) from all biological replicates is indicated below each dataset.

### Use of AI-assisted tools

Generative AI tools, including Claude (Anthropic), Gemini (Google), and ChatGPT (OpenAI), were used to assist with language editing and grammatical refinement of the manuscript. All AI-suggested revisions were critically evaluated and approved by the authors, who take full responsibility for the accuracy and integrity of the final text.

## Acknowledgments

We thank the Drug Discovery Centre supported by the Innovative Support Alliance for Life Science (iSAL), Kyoto University, for imaging facilities used for Western blotting and sister chromatid exchange (SCE) analyses; Yumi Hayashi for technical assistance with molecular cloning; Andrea Ruelas-Gonzalez and Yuya Nishida for experimental support with metaphase-TIF analysis; Minsoo Kim for assistance with Western blot imaging. We also thank all members of the Hayashi laboratory for their support, suggestions, and discussion.

## Author Contributions

P.N. and M.T.H. conceived the study and designed the experiments. P.N. performed most of the experiments and carried out the data analyses presented in this manuscript. M.T.H. contributed to data analysis and interpretation throughout the study, supervised the research, and secured funding. P.N. and M.T.H. wrote the manuscript with input from both authors, and both authors approved the final version.

## Data availability

All data generated or analyzed in this study are available in the article and its Supplementary Information, and from the corresponding author upon reasonable request.

## Ethics approval

This study did not involve human participants or animal models, and therefore did not require ethical approval.

## Abbreviations

AURKB: Aurora kinase B
BLM: Bloom syndrome helicase
BTRR: BLM-TOP3A-RMI1/2 complex
CCR: coiled-coil region
Co-IP: co-immunoprecipitation
D-loop: displacement loop
DDR: DNA damage response
DHBN: Dimerization Helical Bundle in the N-terminal domain
dHJ: double Holliday junction
DSB: double-strand break
dsDNA: double-stranded DNA
FISH: fluorescence in situ hybridization
GEN1: Flap endonuclease GEN homolog 1
HJ: Holliday junction
HR: homologous recombination
HRDC: helicase and RNaseD C-terminal domain
hTERT: human telomerase reverse transcriptase
iDDR: inhibitor of DNA Damage Response
IF-FISH: immunofluorescence-fluorescence in situ hybridization
MAD: mitotic arrest-dependent
meta-TIF: metaphase telomere deprotection-induced foci
MUS81: methyl methanesulfonate (MMS) and ultraviolet (UV) sensitive
NLS: nuclear localization signal
pLDDT: predicted local distance difference tes
PNA: peptide nucleic acid
POT1: protection of telomeres 1
RAP1: repressor and activator protein 1
RMI1/2: RecQ-mediated genome instability protein 1 and 2
RQC: RecQ C-terminal
SCEs: sister chromatid exchanges
shRNA: short hairpin RNA
SLX4: structure-specific endonuclease subunit SLX4
ssDNA: single-stranded DNA
T-loop: Telomere loop
TIF: telomere dysfunction-induced foci
TOP3A: topoisomerase III alpha
TRF1: telomere repeat-binding factor 1
TRF2: telomere repeat-binding factor 2
WRN: Werner syndrome helicase
*γ*-H2AX: phosphorylated histone H2AX at serine 139

