## Supplementary Figures for "The BLM DHBN domain and structure-selective nucleases drive mitotic arrest-dependent telomere deprotection under WRN and TRF2 control"

Supplementary figure 1

a

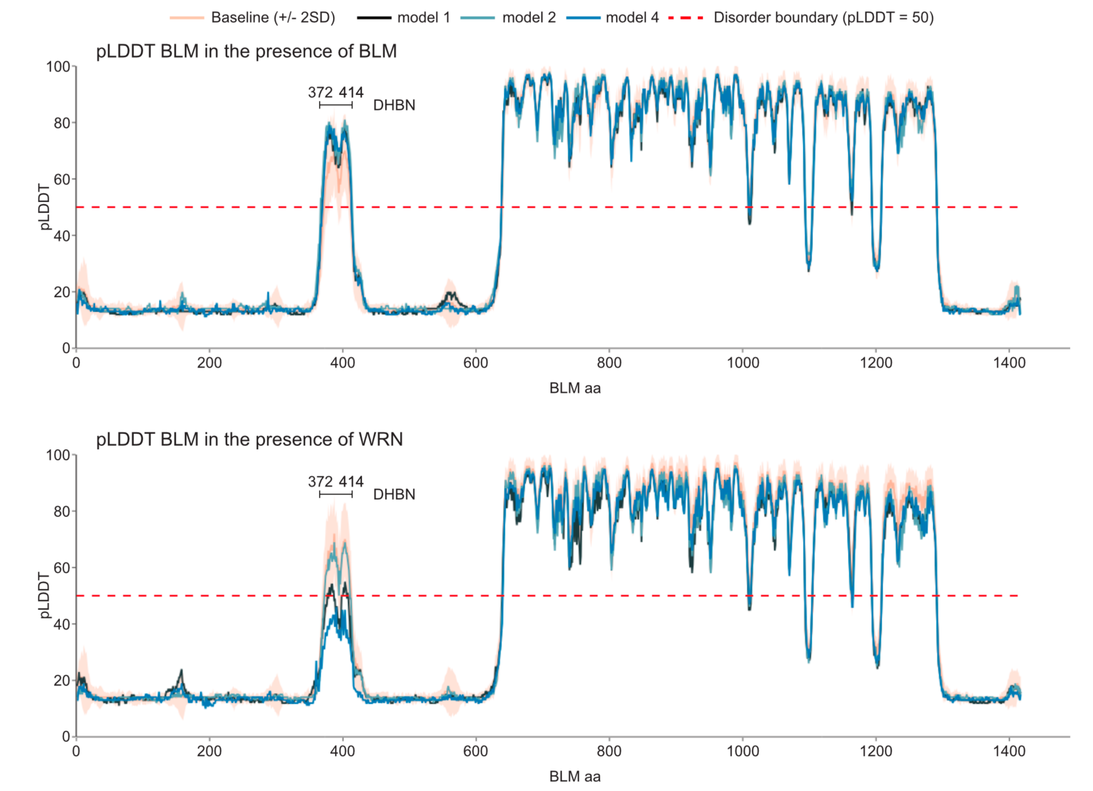


b

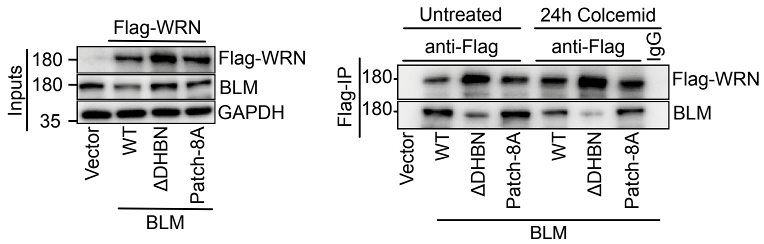


**Supplementary Figure 1. WRN interacts with BLM via the DHBN domain**

**a)** Predicted local distance difference test (pLDDT) scores of BLM modeled as a homodimer (top) and a BLM–WRN heterodimer (bottom). While the DHBN domain exhibits high pLDDT scores in the BLM homodimer, these scores are markedly reduced in the presence of WRN (model 4), suggesting a locally disordered or flexible conformation in the heterodimer context. **b)** Immunoblot analysis confirming the reproducibility of the co-immunoprecipitation (Co-IP) results presented in Figure 1b. Whole-cell extracts and immunoprecipitates were analyzed under the same conditions as in Figure 1b.

**Supplementary figure 2**

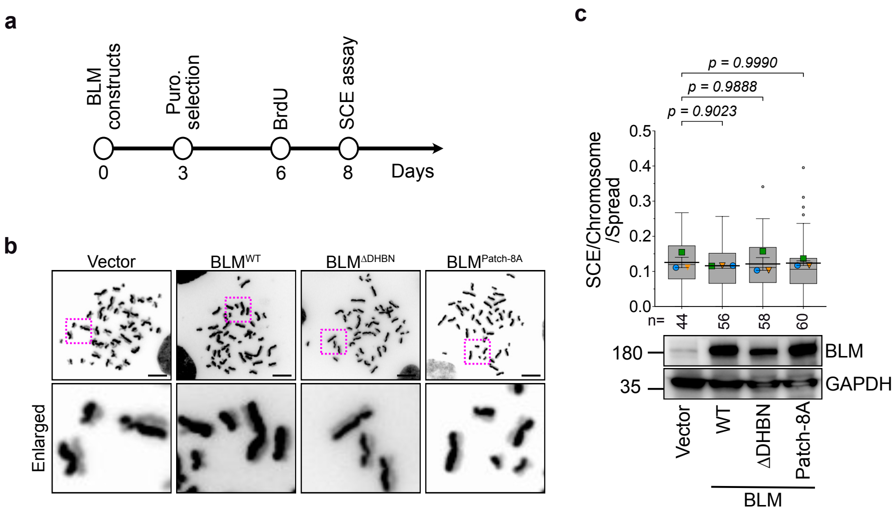


**Supplementary Figure 2. BLM DHBN domain mutants exhibit no dominant negative effect on the anti-SCE activity of endogenous BLM**

**(a)** Schematic of the experimental timeline for the SCE assay in (b, c). **b)** Representative images of sister chromatid exchange (SCE) events in IMR-90 E6E7 hTERT cells expressing the indicated BLM variants. Dashed magenta boxes indicate the regions shown in the enlarged panels. Scale bar, 10 μm. **c)** Top: quantification of SCE events per metaphase spread corresponding to (b). The total number of metaphase spreads analyzed (n) from 3 biological replicates is indicated below each dataset. Data are presented as Tukey box plots of all individual data points, with the mean of each biological replicate superimposed as a distinct colored symbol (circles, squares, and triangles), and the mean ± s.d. of the 3 replicate means is indicated by a horizontal line with error bars. Exact p values are indicated above each comparison; one-way ANOVA followed by Tukey's multiple-comparisons test. Bottom: representative immunoblot analysis confirming the expression levels of the indicated BLM variants at 10 days post-transduction.

**Supplimentary figure 3**

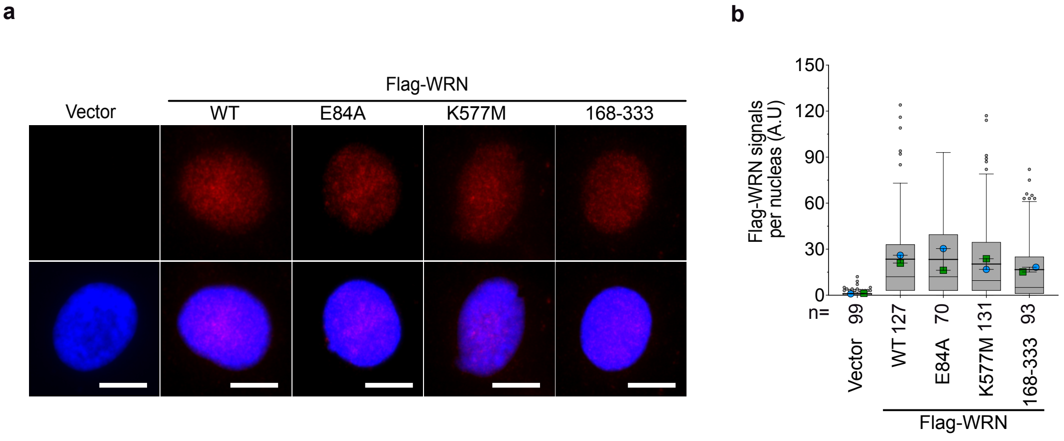


**Supplementary Figure 3. WRN variants localize to the interphase nucleus**

**a)** Representative immunofluorescence images of the Flag-WRN nuclear localization of the indicated WRN variants in interphase cells. IMR-90 E6E7 hTERT cells expressing the indicated 4×Flag-WRN variants were fixed at 8 days post-transduction and immunostained with an anti-Flag antibody (red). DAPI staining (blue) was used to visualize nuclei. Scale bar, 10 µm. **b)** Quantification of Flag-WRN nuclear signal intensity corresponding to (a). The total number of nuclei analyzed (n) from 2 biological replicates is indicated below each dataset. Data are presented as Tukey box plots of all individual data points, with the mean of each biological replicate superimposed as a distinct colored symbol (circles and squares), and the mean of the 2 replicate means is indicated by a horizontal line.

**Supplementary figure 4**

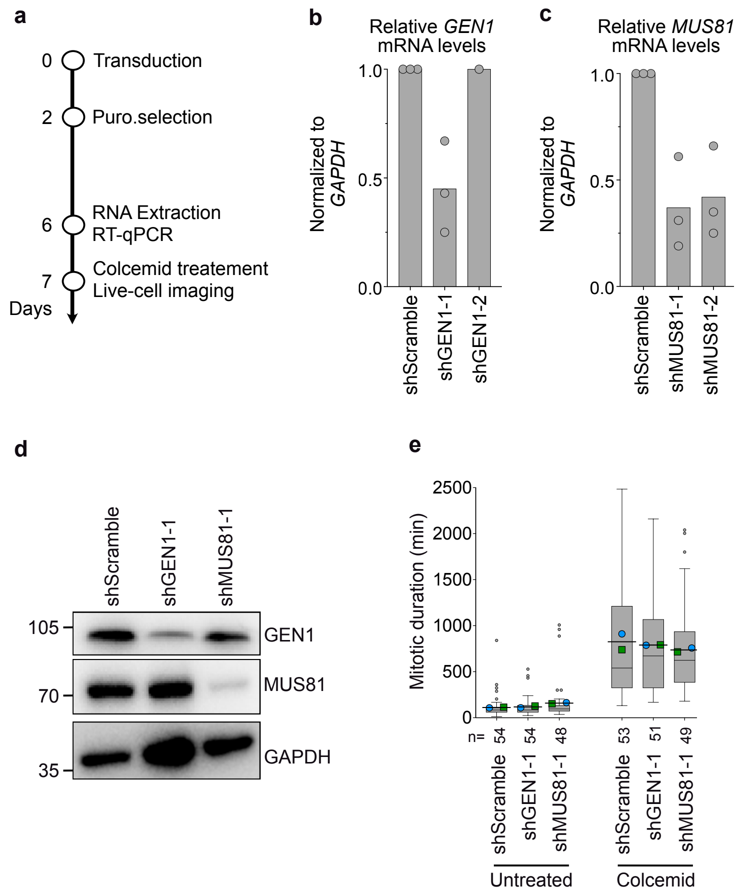


**Supplementary Figure 4. GEN1 and MUS81 knockdowns do not affect mitotic duration upon treatment with a microtubule-depolymerizing agent**

**a)** Schematic of the experimental workflow for lentiviral shRNA-mediated knockdown of GEN1 or MUS81 in IMR-90 E6E7 hTERT cells followed by RT-qPCR and live-cell imaging. **b, c)** RT-qPCR analysis of *GEN1* (b) or *MUS81* (c) mRNA levels in cells transduced with shscramble or two independent shRNAs targeting GEN1 (b) or MUS81 (c). mRNA expression levels were normalized to *GAPDH* and are shown relative to the shScramble control. Data represent the mean of three biological replicates, except for shGEN1-2, for which only one biological replicate was analyzed due to observed cytotoxicity. **d)** Immunoblot analysis confirming depletion of the indicated proteins in cells used for live-cell imaging. **e)** Quantification of mitotic duration in cells corresponding to (d) treated with or without colcemid. The total number of mitotic cells analyzed (n) from 2 biological replicates is indicated below each dataset. Data are presented as Tukey box plots of all individual data points, with the mean of each biological replicate superimposed as a distinct colored symbol (circles and squares), and the mean of the 2 replicate means is indicated by a horizontal line.

**Supplementary figure 5**

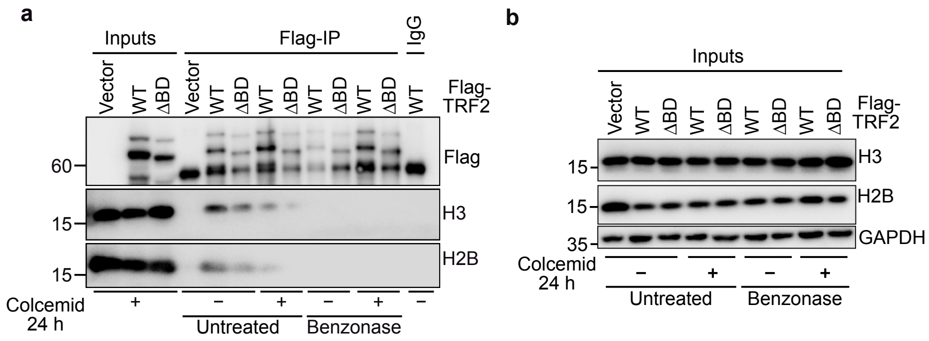


**Supplementary Figure 5. Interaction between the TRF2 basic domain and histones is attenuated during prolonged mitotic arrest**

**a)** Immunoblot analysis confirming the reproducibility of the co-immunoprecipitation (Co-IP) results presented in Figure 6b. Whole-cell extracts and immunoprecipitates were analyzed under the same conditions as in Figure 6b to confirm the robustness of the TRF2–histone interaction and its attenuation during mitotic arrest. **b)** immunoblot analysis demonstrating that histone H2B and H3 protein levels remain unchanged following colcemid or Benzonase treatment.

**Supplementary figure 6**

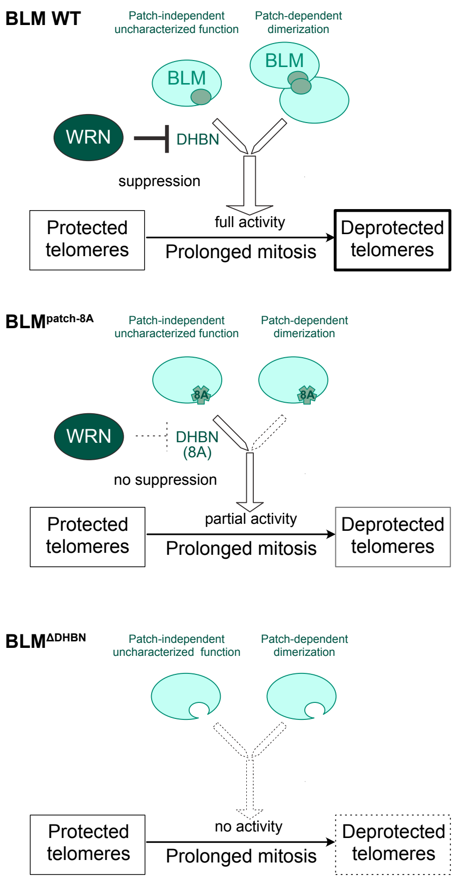


**Supplementary Figure 6. Proposed model for WRN-mediated suppression of BLM-dependent MAD telomere deprotection.**

BLM promotes MAD telomere deprotection through its DHBN domain, which harbors two distinct functions: hydrophobic patch-dependent dimerization/multimerization and a patch-independent uncharacterized function. WRN directly associates with BLM via the DHBN domain and restrains these activities, thereby limiting MAD telomere deprotection. In cells expressing BLM$^{Patch\text{-}8A}$, the mutant fails to dimerize/multimerize but retains the uncharacterized function, which is sufficient to support partial MAD telomere deprotection. WRN continues to interact with BLM$^{Patch\text{-}8A}$; however, the Patch-8A mutations render BLM$^{Patch\text{-}8A}$ insensitive to WRN-mediated suppression of this uncharacterized function. In cells expressing BLM$^{\Delta DHBN}$, BLM completely loses both functions and is therefore unable to promote MAD telomere deprotection.

**Supplementary figure 7**

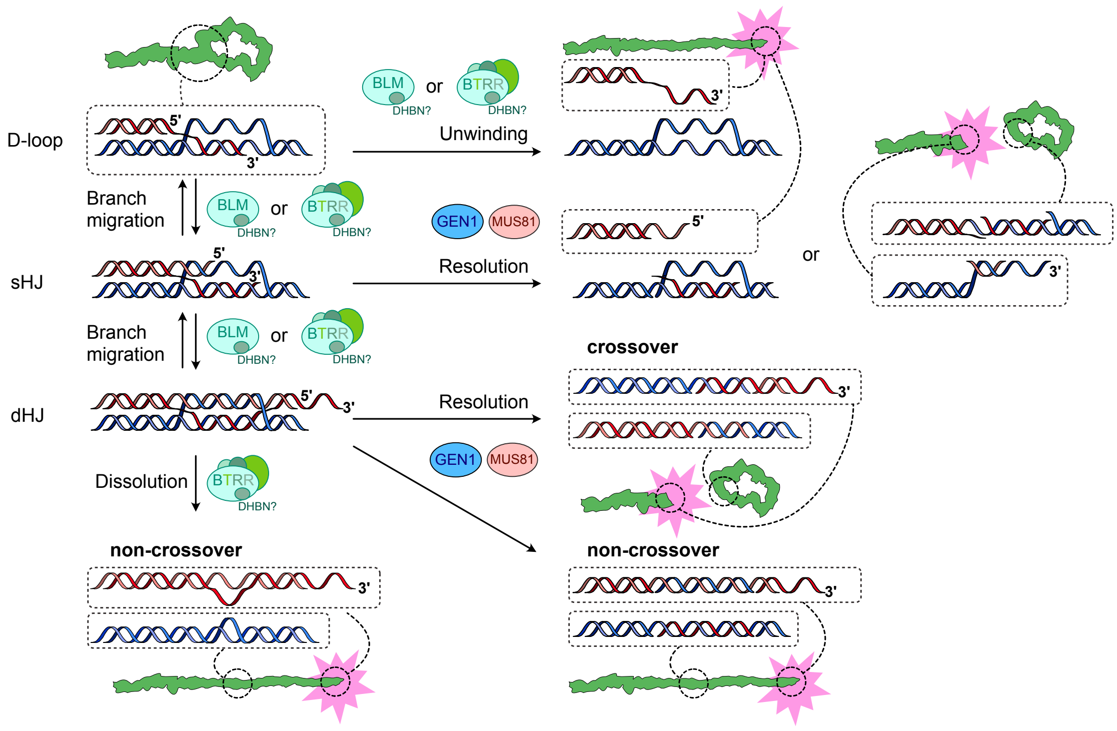


**Supplementary Figure 7. Proposed model for MAD telomere deprotection.**

Prolonged mitotic arrest creates a permissive state for BLM recruitment to the T-loop by weakening TRF2-dependent telomere protective mechanisms. Through a DHBN domain-dependent activity, BLM promotes remodeling of the T-loop structure, potentially generating single Holliday junction (sHJ) or double Holliday junction (dHJ) intermediates. Formation of these intermediates may be facilitated by BLM oligomerization and/or branch migration activity. The resulting recombination-like DNA structures then become substrates for either processing by the BTRR complex through dissolution or nucleolytic resolution by the structure-selective endonucleases MUS81 and GEN1. Processing of these intermediates disrupts the protective T-loop architecture, leading to telomere deprotection and activation of the telomeric DNA damage response. In this model, the DHBN domain serves as a critical regulatory element that enables BLM-dependent generation of telomeric recombination intermediates, thereby promoting MAD telomere deprotection.
