## Supplementary Material for "The BLM DHBN domain and structure-selective nucleases drive mitotic arrest-dependent telomere deprotection under WRN and TRF2 control"

Original blots (pages 1–8) for immunoblot analysis performed in this study.

Membranes were routinely cut before primary antibody incubation. The top and bottm edges of each membrane are indicated by magenta arrows.

**Original blots 1**

### Blots in Fig. 1b

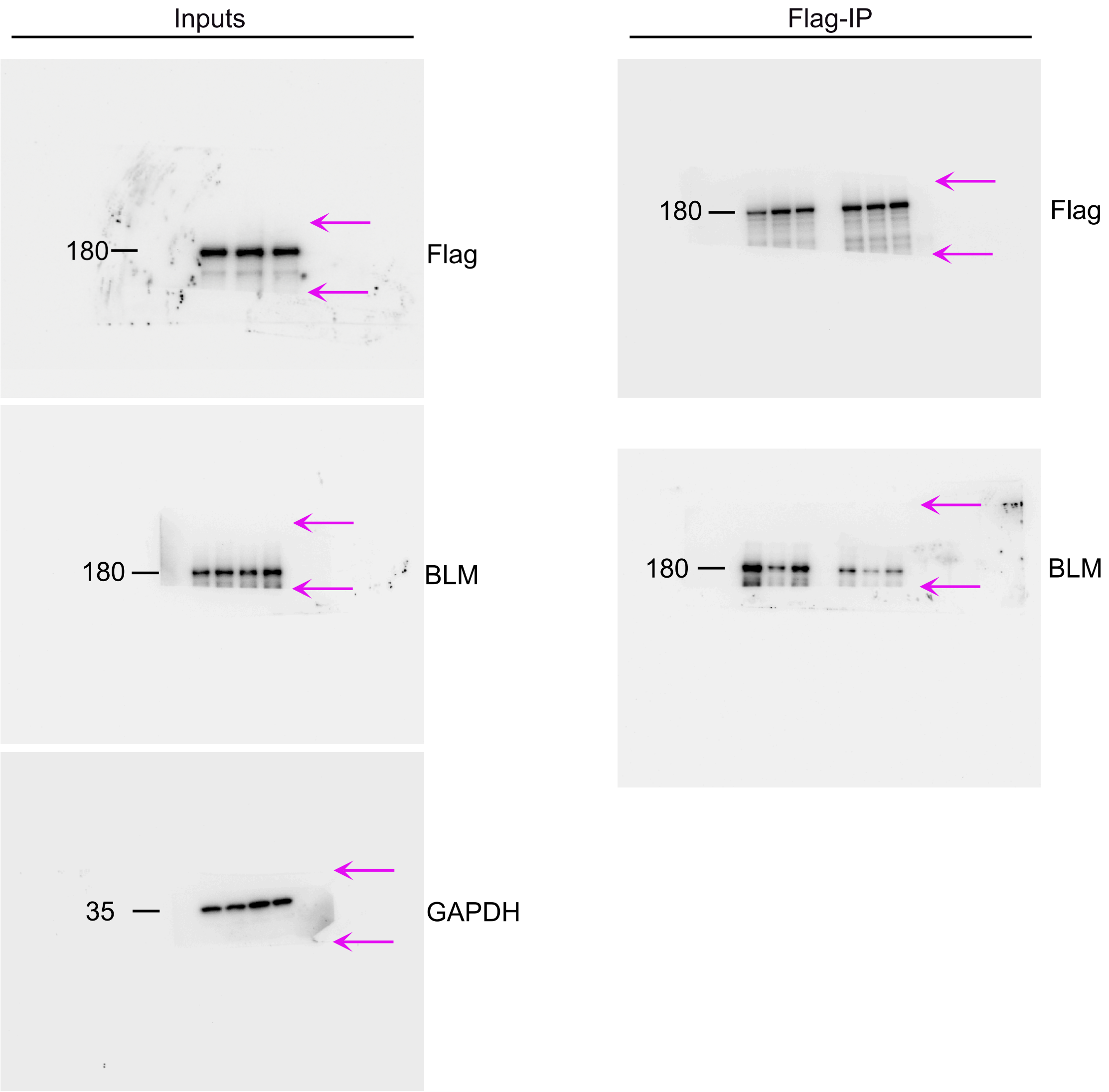


### Blots in Fig. 1c

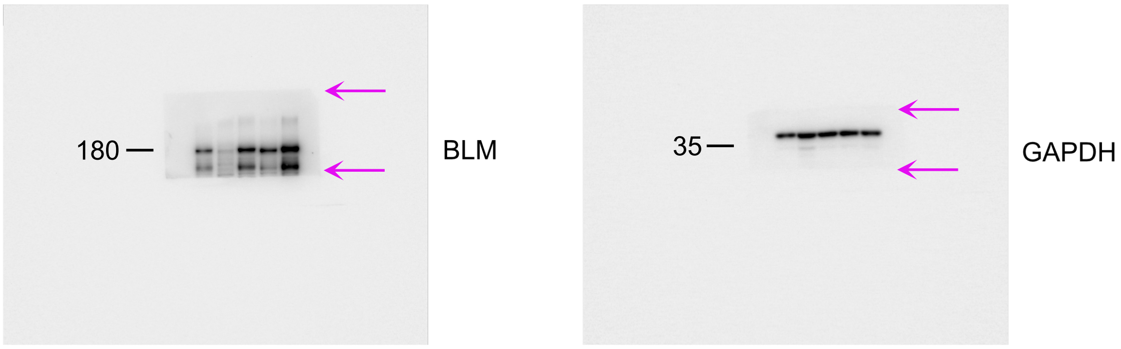

**Original blots 2**

### Blots in Fig. 2b

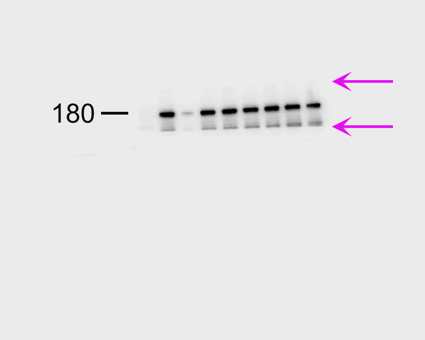


BLM

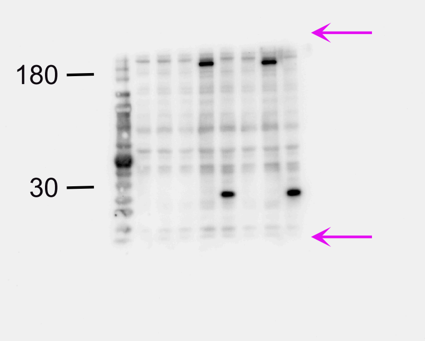


Flag

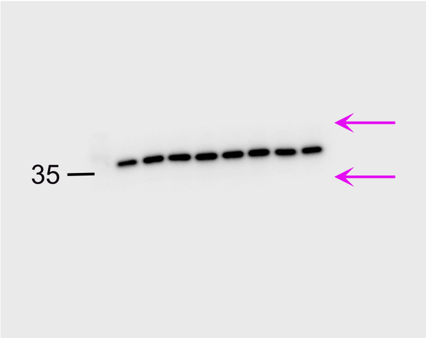


GAPDH

**Original blots 3**

### Blots in Fig. 3c

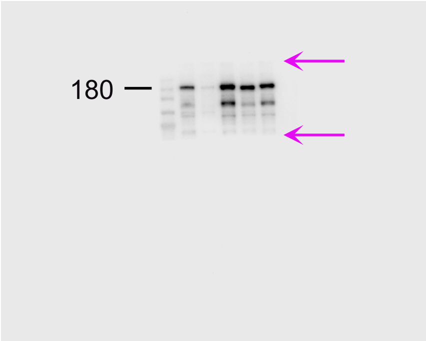


BLM

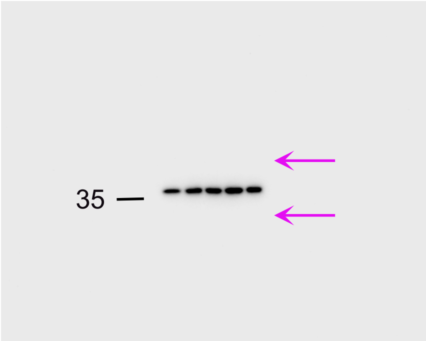


GAPDH

### Blots in Fig. 3f

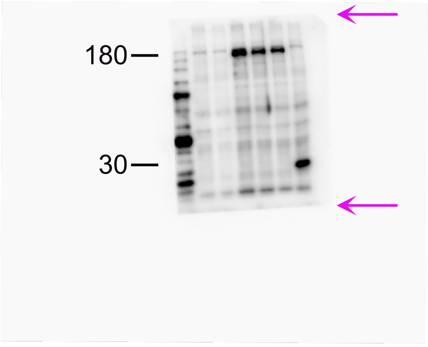


Flag

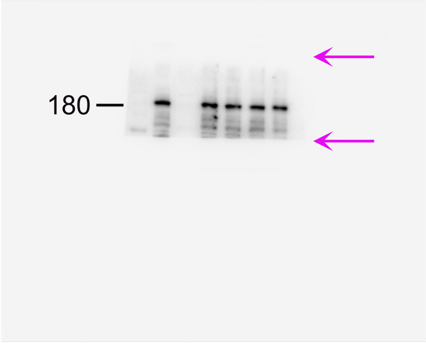


BLM

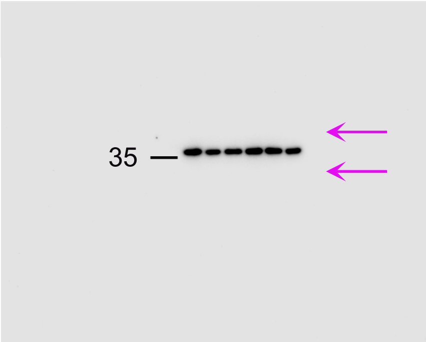


GAPDH

**Original blots 4**

#### Blots in Fig. 4b

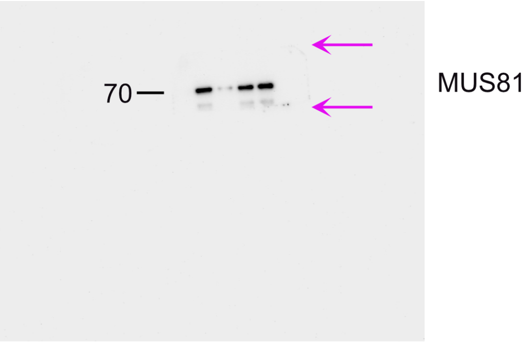


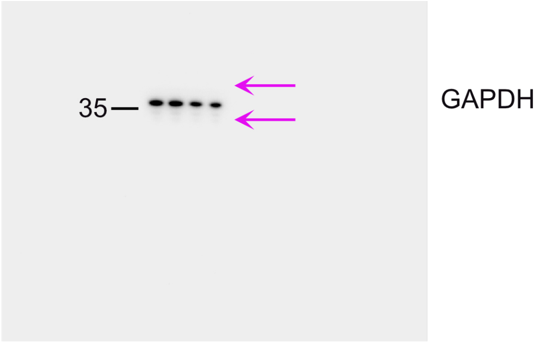


#### Blots in Fig. 5b

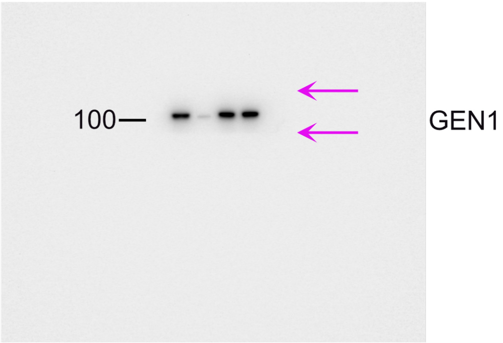


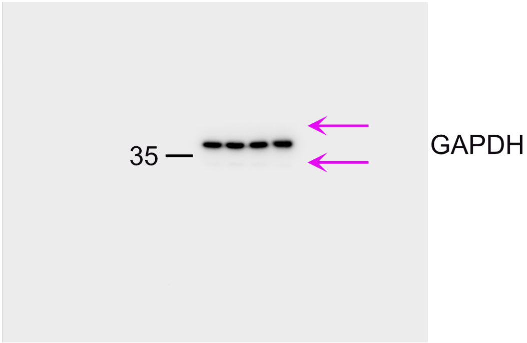

**Original blots 5**

#### Blots in Fig. 6b

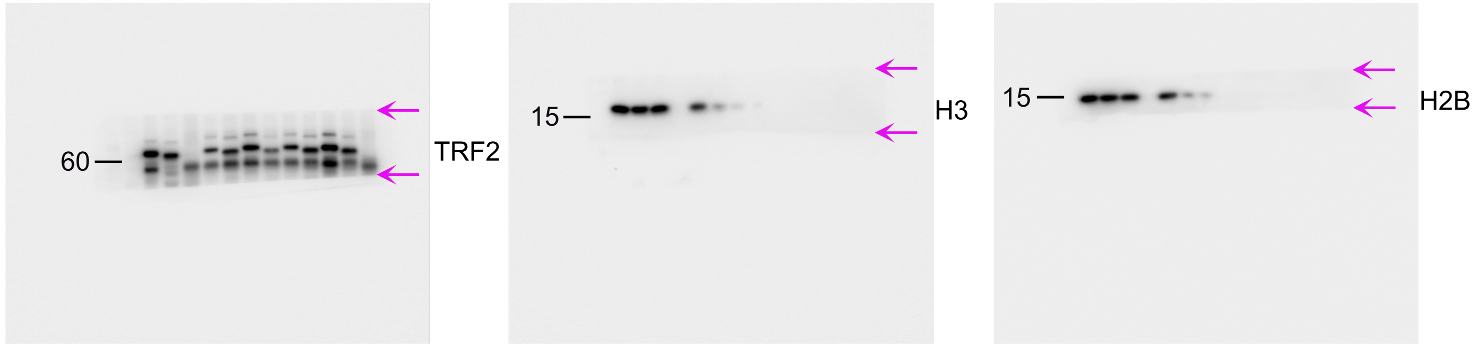


#### Blots in Fig. 6c

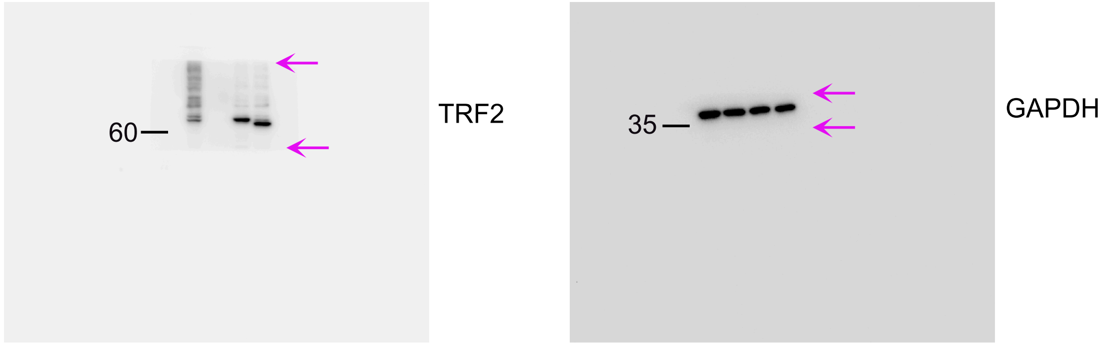


#### Blots in Fig. 6g

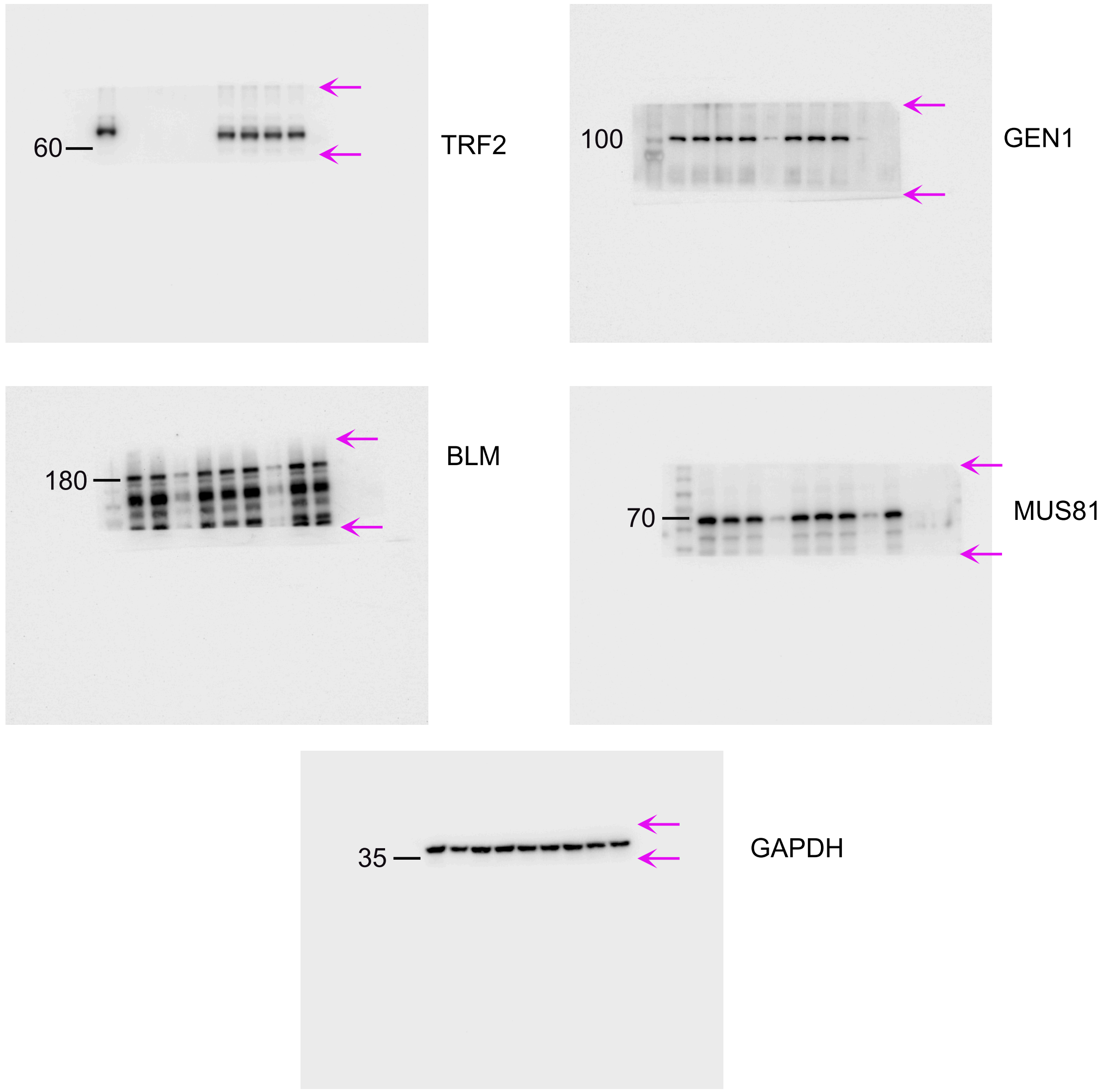

**Original blots 6**

### Blots in Fig. S1b

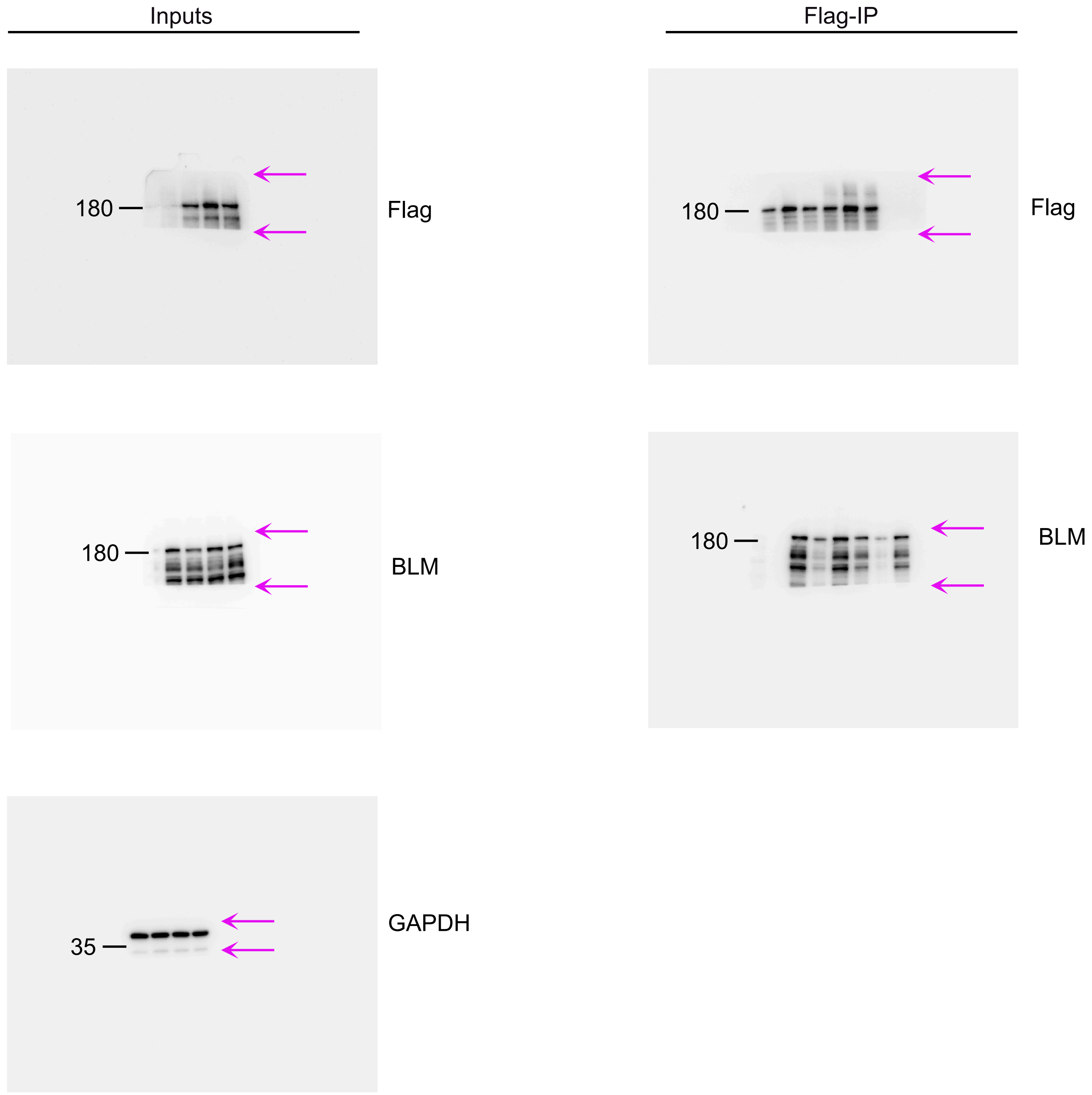

**Original blots 7**

#### Blots in Fig. S2c

BLM

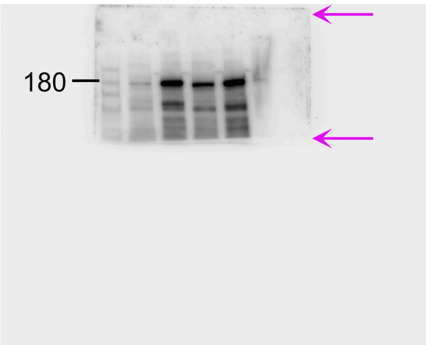


GAPDH

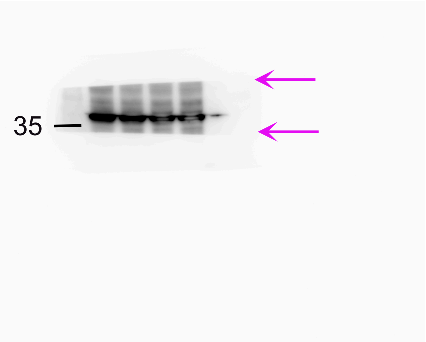


#### Blots in Fig. S4b

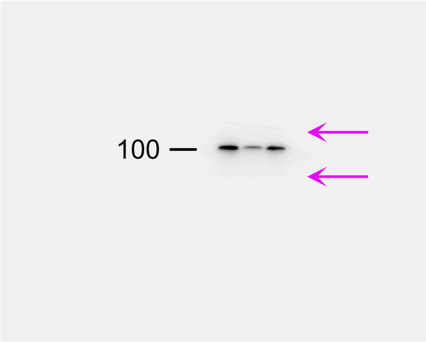


GEN1

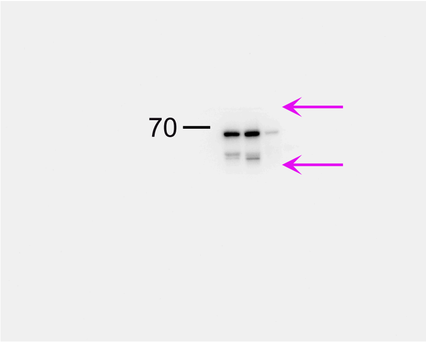


MUS81

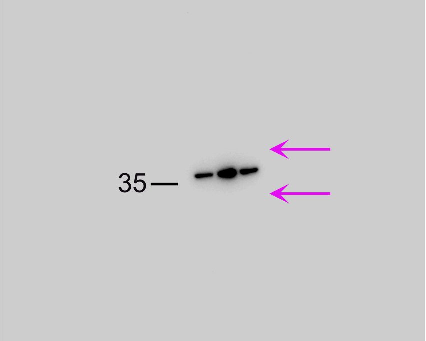


GAPDH

**Original blots 8**

#### Blots in Fig. S5a

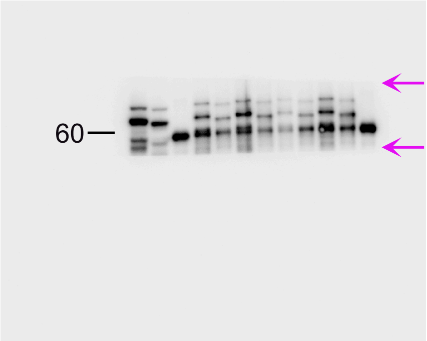


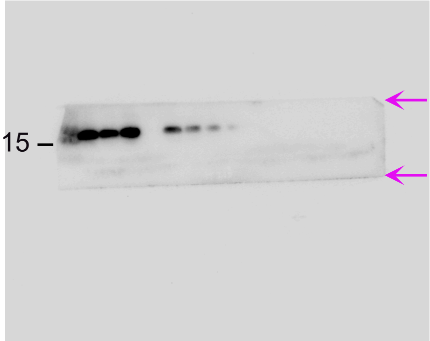


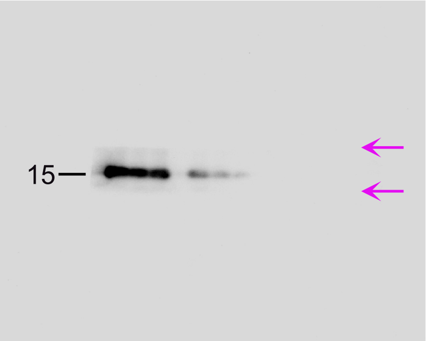


#### Blots in Fig. S5b

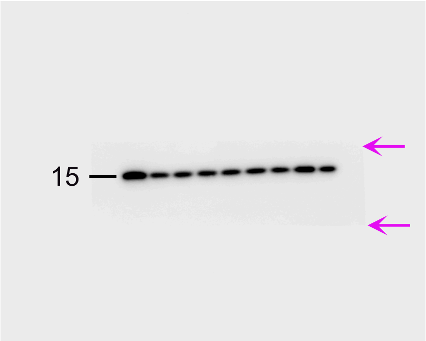


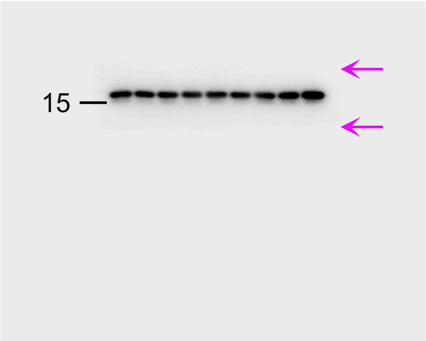


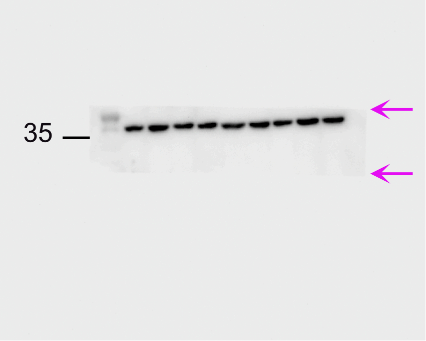
